# Epitope-Driven Polyfunctional Divergence of SARS-CoV-2 RBD Antibodies

**DOI:** 10.1101/2025.08.26.672080

**Authors:** Pierre Rosenbaum, Cyril Planchais, Timothée Bruel, Maxime Beretta, Isabelle Staropoli, William-Henry Bolland, Florence Guivel-Benhassine, Delphine Planas, French COVID Cohort Study Group, CORSER Study Group, Olivier Schwartz, Hugo Mouquet

## Abstract

SARS-CoV-2 antibodies targeting the receptor binding domain (RBD) of the spike protein can potently neutralize infection and exert additional antiviral functions. Here, we characterize the functional profiles of human RBD-specific memory B-cell antibodies elicited by ancestral SARS-CoV-2 infection. While SARS-CoV2 neutralizing antibodies, mainly class 1 and 3 anti-RBD antibodies, often lose binding and neutralizing capacities against Omicron variants, most non-neutralizers remain broadly reactive. Despite their restricted cross-reactivity, neutralizing antibodies mediate Fc-effector functions including antibody-dependent cellular cytotoxicity, phagocytosis, and complement deposition. Most neutralizers also enhance binding of antibodies that target the SARS-CoV-2 spike fusion peptide *via* receptor-mimetic allostery. In contrast, broadly reactive non-neutralizers fail to trigger phagocytic or allosteric effects. Thus, viral escape compromises antibody neutralizing as well as other antiviral activities, leaving broadly reactive, non-polyfunctional antibodies that exert minimal immune pressure less effective at controlling infection.

## INTRODUCTION

Humoral immune responses to SARS-CoV-2, most notably neutralizing antibodies, are essential for protecting vaccinated and convalescent individuals from (re-)infection^1–3^. SARS-CoV-2 neutralizing antibodies target multiple epitopes on three major regions of the viral spike glycoprotein including the receptor-binding domain (RBD), N-terminal domain (NTD) and S2 stalk^4,5^. Anti-RBD neutralizers belong to six different classes, depending on their epitope location - either within (class 1 and 2) or outside (class 3 to 6) the receptor-binding motif (RBM)^6–8^. They are major contributors to seroneutralizing activity^9^, and many have been engineered as monoclonal antibodies for both prophylactic and therapeutic interventions against COVID-19^10,11^. However, the antigenic drift resulting from the continuous evolution of SARS-CoV-2 has severely compromised the effectiveness of both antibodies pre-existing in immune donors and those used in clinics^12–17^. Novel anti-RBD neutralizing antibodies that were more resilient to viral escape - such as SA55, VIR-7229 and VYD2311 - were advanced into clinical development^18–20^, but only SA55 remains potently active against the most recent post-JN.1 variants^21^. Apart from neutralization, SARS-CoV-2 spike-specific antibodies can harbor Fc-effector functions such as antibody-dependent cell cytotoxicity (ADCC), antibody-dependent cell phagocytosis (ADCP) and complement dependent cytotoxicity (CDC), which could contribute to SARS-CoV-2 infection control^22–31^. Infection and vaccination elicit SARS-CoV-2 antibodies with Fc-dependent effector functions that persist over time^30–32^. Although their exact roles remain to be fully elucidated, these functions may indeed be critical for viral clearance, as exemplified by studies on class 3 antibody S309/Sotrovimab^33,34^. Certain anti-RBD antibodies also exert allosteric effects by inducing conformational changes in the spike protein that mimic the angiotensin converting enzyme 2 (ACE2) interaction, thereby enhancing the potency of non-RBD neutralizing antibodies^35–37^.

We previously uncovered a potential dichotomy in SARS-CoV-2 antibody-mediated antiviral functions according to spike epitope specificity, with potent Fc-dependent effectors mainly targeting the S2 subunit and NTD; however, only a small subset of anti-RBD monoclonal antibodies were characterized in that study^38^. Here, to gain deeper insight into Fc-effector antiviral mechanisms of anti-RBD antibodies, we generated a panel of human RBD-specific monoclonal antibodies cloned from memory B cells of individuals who recovered from ancestral SARS-CoV-2 infection. We characterized their reactivity and epitope profiles, as well as their antiviral activities including neutralization, ACE2-mimetic effects, and Fc-dependent effector functions. We showed that Omicron lineages preferentially evade polyfunctional class 1 anti-RBD neutralizers, reducing not only neutralization but also associated Fc-effector and ACE2-mimetic allosteric activities, whereas the remaining broadly reactive non-neutralizing anti-RBD antibodies have substantially diminished functional capacity.

## RESULTS

### Human anti-RBD memory B-cell antibody profiles

To select ancestral SARS-CoV-2-exposed convalescent donors (n=42 total) for single B-cell antibody cloning, we first evaluated their IgG seroreactivity to recombinant Wuhan, Beta (β) and Delta (δ) SARS-CoV-2 RBD proteins by ELISA (**Figs. 1A** and **S1A**). Peripheral blood IgG^+^ and IgA^+^ memory B cells from five donors with high serum anti-RBD antibody levels were stained with fluorescently labeled β and δ RBD protein baits for flow cytometry phenotyping and single-cell sorting (**Figs. 1B** and **S1B**). RBD-dual reactive class-switched memory B cells (RBD^βδ+^) were detected at similar frequencies among donors (0.11±0.02%) and mainly displayed a resting memory phenotype (CD19^+^CD27^+^CD21^+^) (**Figs. 1B**, **1C**, **S1C** and **S1D**). From single RBD^βδ+^-captured B cells, we produced 66 unique recombinant IgG1 monoclonal antibodies by expression cloning^39^ (originally, 49 IgG and 17 IgA as native isotype) (**Table S1**). Among them, 84% (n=56) were specific based on high-affinity ELISA binding to Wuhan RBD, and as expected, the vast majority also recognized both β and δ variant RBD proteins (84%, n=47) (**Figs. 1D** and **1E**). Subsequent analyses were restricted to dual-reactive (β and δ) anti-RBD antibodies and excluded Wuhan-monoreactive antibodies (n = 9). To assess their cross-reactive potential against viral variants, we first performed ELISA and flow cytometry binding analyses against 23 different SARS-CoV-2 RBD and/or S proteins (Alpha to Omicron BA.2.86). About half of them (45%) showed pan-reactivity against pre- and post-omicron variants despite being derived from memory B cells elicited by the ancestral strain (**Figs. 2A**, **S2A** and **Table S2**). Some anti-RBD^βδ^ antibodies (17%, n=8) strongly cross-reacted with the SARS-CoV-1 tri-S protein, but not with those of 229E, MERS-CoV, or HKU1 (**Fig. S2B**). Based on RBD-ELISA competition assay using reference antibodies as potential competitors - REGN10987^40^, CB6^41^, LY-CoV555^42^, CT-P59^43^, COV2-2130^44^, ADG-2^45^, Ly-CoV1404^13^ and S309^46^ - we categorized the anti-RBD antibodies as class 1 (34%), class 2 (8.5%), class 3 (10.6%) or unclassified (46.8%) (**Fig. 2B**). Antibodies with restricted binding profiles against RBD variants primarily belonged to class 1 and 2 (81% and 75%, respectively), whereas those with broader reactivity spectra were more frequently found in class 3 and unclassified groups (60% and 64%, respectively) (**Figs. 2C, 2D** and **S2C**).

**Figure 1.**
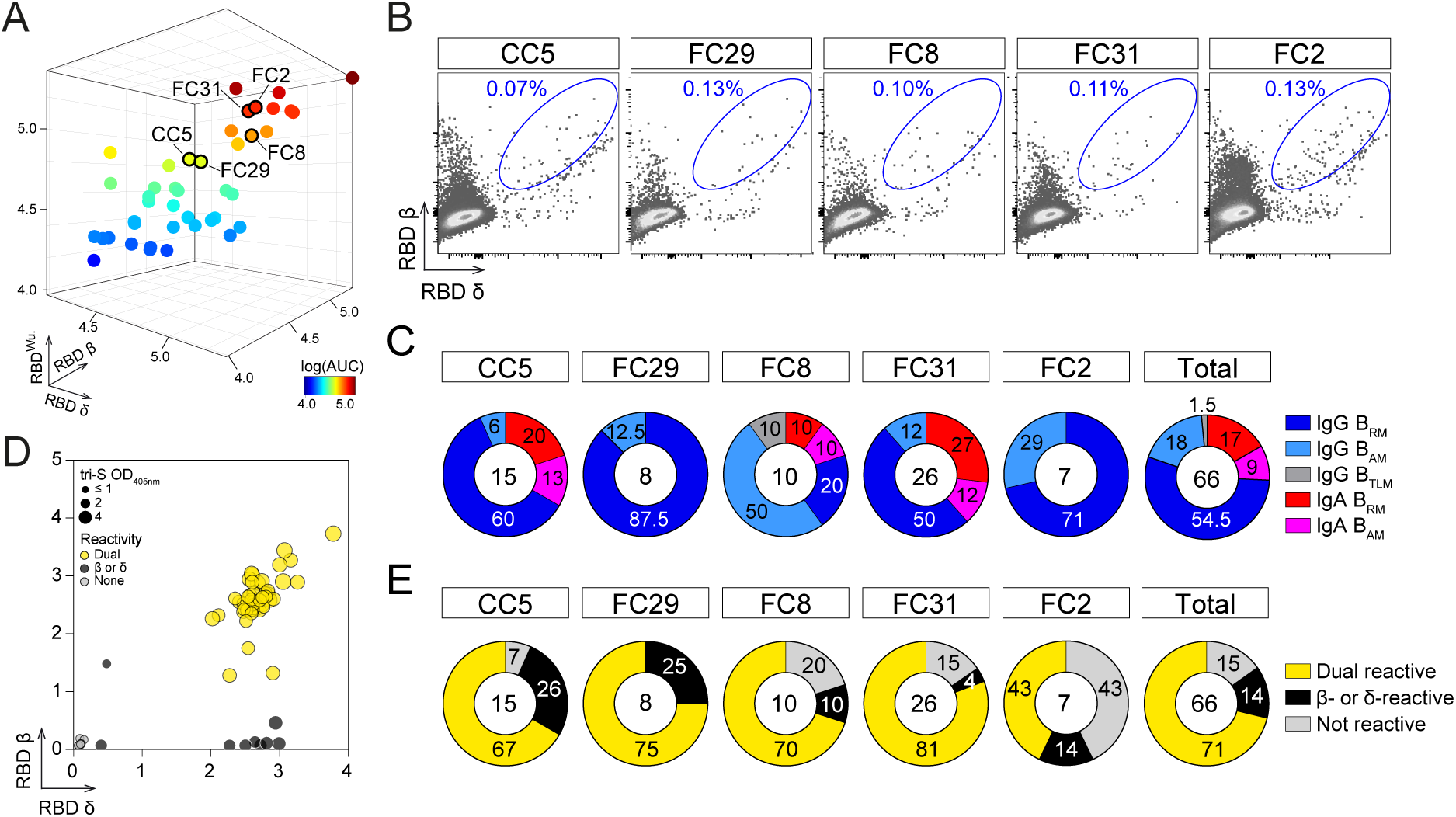
Human SARS-CoV-2 RBD-specific memory B-cell antibodies. (**A**) 3D plot comparing the IgG seroreactivity of COVID-19 convalescent individuals (n=42) against purified Wuhan (Wu.), Beta (β), and Delta (δ) RBD proteins. Labeled, circled dots indicate donors selected for single memory B-cell flow cytometry sorting. Dots are color-coded according to their ELISA reactivity measured as areas under curve (AUC). (**B**) Flow-cytometric plots showing circulating blood RBD-dual reactive (RBD^βδ+^) IgG^+^ and IgA^+^ memory B cells (gated as indicated in Fig. S1B) from the 5 selected convalescent donors. (**C**) Donut charts comparing the phenotypic distribution of IgG^+^ and IgA^+^ memory B cells from which anti-RBD antibodies have been cloned (total cloned antibodies *per* donor are indicated in the center of the charts: B_RM_ (resting memory, CD27^+^CD21^+^), B_AM_ (activated memory, CD27^+^CD21^−^), and B_TLM_ (tissue-like memory, CD27^−^CD21^−^). Value at the center of each chart represents the total amount of antibodies *per* donor. (**D**) Bubble plot comparing the ELISA reactivity of human RBD^βδ+^-captured memory B-cell antibodies against RBD Delta (δ, x axis), RBD Beta (β, y axis) and tri-S Wuhan (bubble size). OD, optical dentities. Antibodies were categorized as dual (yellow; OD > 1 for both RBD β and RBD δ), variant-specific (black; OD > 1 for RBD β or RBD δ), and not reactive (gray; OD < 1 for all RBDs). (**E**) Donut charts comparing the distribution of anti-RBD antibody profiles based on the ELISA reactivities measured in Fig.1D. Total cloned antibodies *per* donor are indicated in the center of the charts.

**Figure 2.**
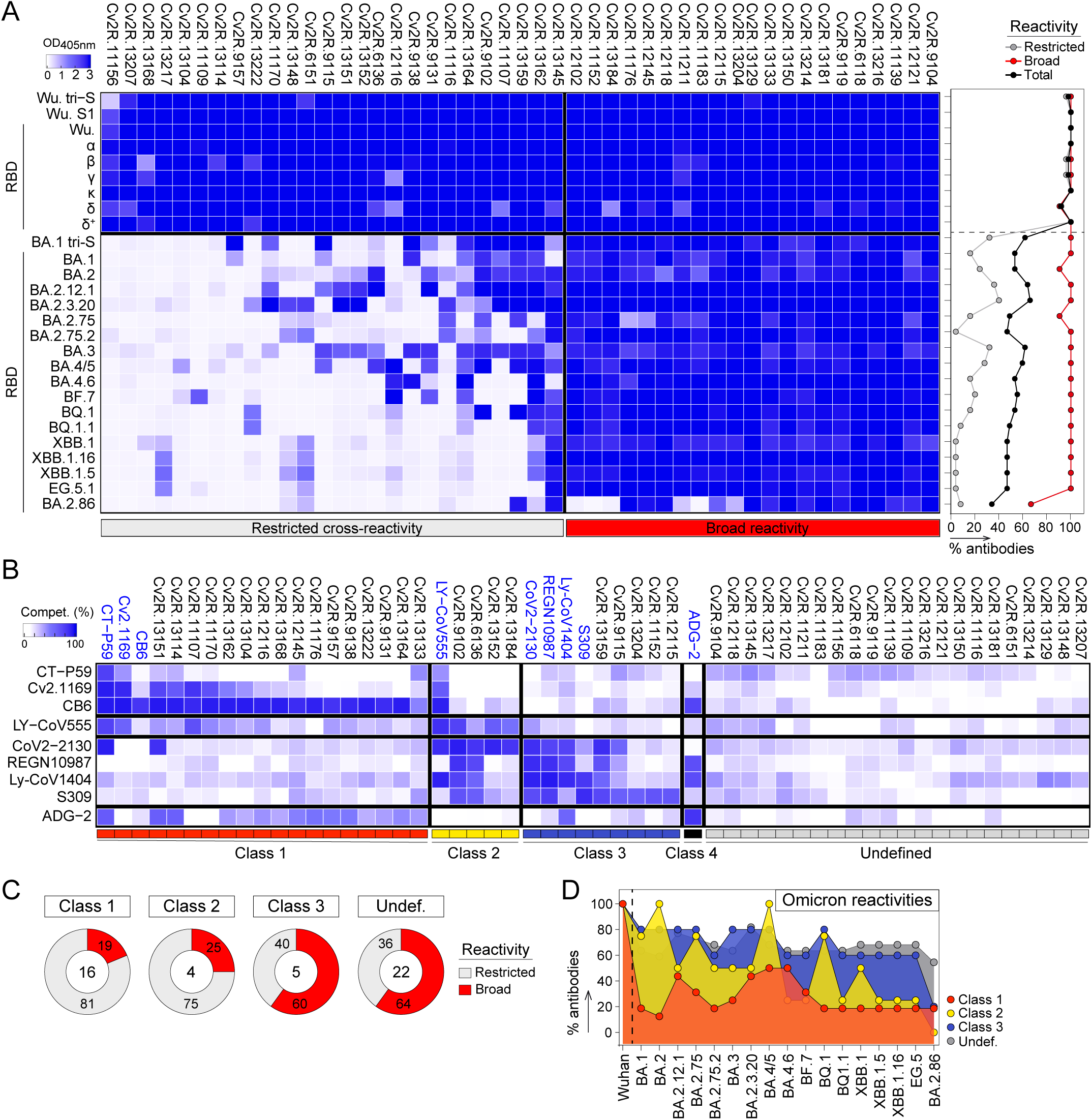
Reactivity profiles of human anti-RBD antibodies. (B) Heatmap (left) comparing the ELISA binding profiles of human anti-RBD antibodies (n = 47) against purified recombinant SARS-CoV-2 Wuhan (Wu.) tri-S, S1 and RBD proteins from various viral variants. Cells are color-coded according to mean OD values obtained with antibodies at 1 µg/ml. Antibodies were categorized as either broadly reactive (red) or with a more restricted cross-reactivity (grey). Graph (right) showing the proportion of antibodies in each category). (**B**) Heatmap showing competition for Wuhan RBD binding of anti-RBD antibodies in the presence of reference antibodies as potential competitors: CT-P59, Cv2.1169, and CB6 (class 1); LyCoV555 (class 2); REGN10987, LyCoV1404, CoV2-2130 and S309 (class 3); ADG-2 (class 4). Cells are color-coded based on the binding inhibition relative to reactivity in absence of potential competitor (% bind.); dark blue corresponds to high binding inhibition, while white indicates no competition. Mean of duplicates values are shown. (**C**) Donut charts comparing the frequency of broadly reactive (red) and restricted cross-reactive (gray) antibodies for each RBD class identified in (B) (1, 2, 3 and undefined). Total number of antibodies in each group is indicated in the center of the charts. (**D**) Graph comparing the reactivity profiles of anti-RBD antibodies against SARS-CoV-2 RBD proteins from Wuhan and Omicron variants according to their class as defined in (B). Antibodies were considered reactive when OD > 2, and their proportion (%) is depicted (y axis).

### Neutralizing and Fc-dependent effector activities of human anti-RBD antibodies

We next evaluated the *in vitro* neutralizing capacity of anti-RBD antibodies against authentic live viruses using the S-Fuse assay^47^. About half of them (n=20) showed significant neutralizing activity against at least two variants in a pre-Omicron 5-viruses panel (D614G, α, β, γ, and δ), with a quarter being active against all five viral strains (**Fig. 3A** and **Table S3**). Most cross-neutralizers belonged to class 1 and 3 RBD antibodies (**Fig. S3A**). Comparable results were observed using a virus like particles (VLP) reporter cell-based assay and an RBD-ACE-2 competitive ELISA^39^, with blocking activity in the latter assay mainly restricted to class 1 antibodies, as anticipated (**Figs. 3A**, **S3A**-**C** and **Table S3**). We further examined the breadth of the six most potent neutralizers (5 class 1 and a class 3) against 8 Omicron variants in the S-Fuse assay (BA.1 to XBB.1.5). As previously observed for other class I such as Cv2.1169^48^, all lost their activity post-BA.2 excepted Cv2R.13162, which still neutralized SARS-CoV-2 variants up to XBB.1.5 (**Figs. S4A-D**). Noteworthy, antibodies with broad reactivity profiles on pre- and Omicron RBD proteins were more frequently found among non-(cross)neutralizers as compared to cross-neutralizing antibodies (70% *vs* 10%, p=0.0001) (**Figs. 3A** and **3B**). To further investigate the antiviral properties of human SARS-CoV-2 RBD-specific antibodies, we next assessed their *in vitro* capacity to mediate ADCC, ADCP and complement deposition (CD) as a proxy for CDC. Most anti-RBD antibodies induced both ADCC and CD to varying degrees, irrespective of whether their RBD reactivity spectra were narrowed or broad (**Figs. 3C**, **3D** and **Table S3**). In contrast, only few broadly RBD-reactive antibodies promoted ADCP, compared with antibodies exhibiting limited cross-reactivity (9.5 % *vs* 54%, p=0.002) (**Figs. 3C**, **3D** and **Table S3**). Moreover, potent ADCP inducers were more frequently found among class 1 RBD antibodies, the majority of which neutralized only pre-Omicron variants (**Fig. S3D** and **Table S3**). These data show that neutralizing anti-RBD antibodies can also serve as broad Fc-dependent effectors, while broadly RBD-reactive antibodies are non-neutralizing, less polyfunctional, but can still trigger ADCC and CD.

**Figure 3.**
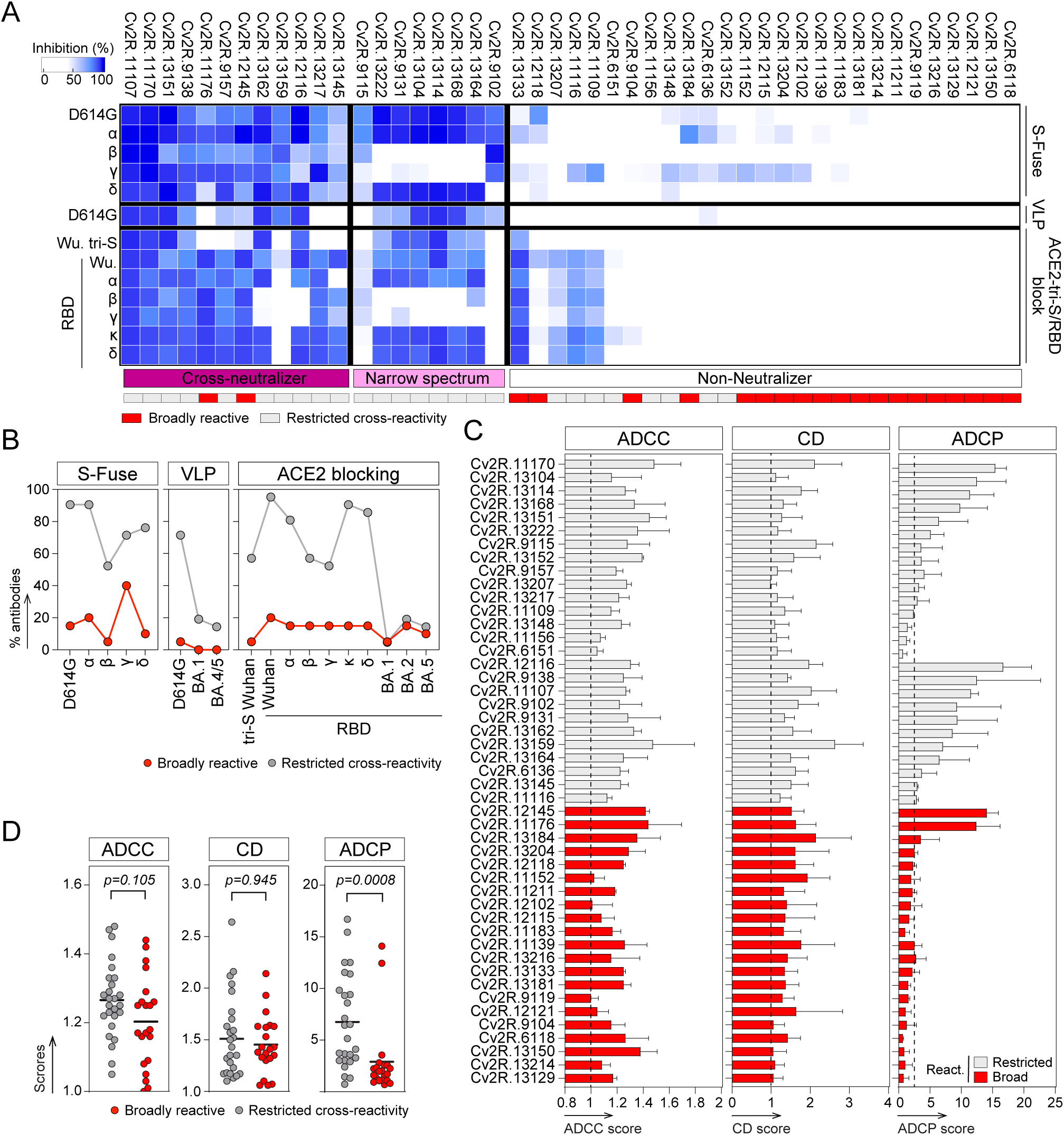
Neutralizing and Fc-dependent effector activities of human anti-RBD antibodies. (**A**) Heatmap comparing the *in vitro* neutralization activity of human anti-RBD antibodies against SARS-CoV-2 D614G, Alpha (α), Beta (β), Gamma (γ) and Delta (δ) viruses using the S-fuse, virus-like particles (VLP) neutralization assays, and ACE2-tri-S or -RBD blocking ELISA assay. Dark blue colors indicate strongest inhibition percentages, while white corresponds to no inhibition. Mean of duplicate values are shown. Antibodies were categorized as either cross-neutralizing (dark pink), neutralizing (pink) or weak or non-neutralizing (white). (**B**) Graph comparing the proportion of anti-RBD antibodies active in the S-Fuse, VLP, and ACE2/tri-S|RBD assays according to their reactivity profiles as defined in Fig. 2A. Antibodies were considered active when inhibition/neutralization % > 50%. (**C**) Bar graphs comparing the ADCC, CD, and ADCP activities of anti-RBD antibodies (n=47). Bars are colored-coded based on antibody reactivity profiles. Data represents the mean± SD of duplicates from two independent experiments. Dotted lines represent positivity threshold based on the negative control. (**D**) Scatter plots comparing the ADCC, CD and ADCP scores of anti-RBD antibodies according to their reactivity profiles (broad, n=21 and restricted, n=26). Groups were compared using Mann-Whitney’s test.

### ACE2-mimetic property of human anti-RBD antibodies

Beyond direct antiviral effects, anti-RBD antibodies can allosterically or cooperatively enhance the activity of other non-RBD antibodies^49–53^. Some of them increase S2 epitope accessibility, thereby boosting the neutralizing capacity of anti-S2 antibodies, including those targeting the FP^36,37^. Considering the potential *in vivo* relevance of cross-epitope antibody cooperativity and synergy, we next examined whether anti-RBD antibodies could facilitate exposure of the FP epitope. To this end, we produced four novel human anti-FP antibodies cloned from FP-captured memory B cells of two selected convalescent donors with high serum anti-FP IgG levels (**Figs. S5A-D**). Three of them (Cv2F.14107, Cv2F.14125, and Cv2F.14132) were clonal variants from the same B-cell lineage, whereas the other (Cv2F.4147) was a unique clone (**Table S1**). Despite their broad reactivity across coronaviruses, including SARS-CoV-2 variants, they showed little or no neutralizing activity against SARS-CoV-2, unlike previously reported anti-FP antibodies 76E1, CoV4462 and C77G12^36,37,54^ (**Figs. 4A** and **S5E-G**). Following soluble ACE2 interaction, Cv2F.14107, but not Cv2F.4147, showed increased reactivity to cell-expressed δ spike proteins (**Fig. 4B**), and was therefore used as a probe in the flow cytometry binding assay to reveal allosteric effects of anti-RBD antibodies. Most antibodies that enhanced Cv2F.14107 binding to the SARS-CoV-2 spike had narrow RBD cross-reactivity (81% *vs* 14% for broad, p<0.0001) and, consequently, mainly belonged to class 1 and neutralizer groups (**Figs. 4C**, **4D** and **S6A**). Nevertheless, some non-neutralizing (Cv2R.13148, Cv2R.6151) and broadly reactive (Cv2R.12145, Cv2R.11176) antibodies also triggered FP epitope exposure, indicating that this cooperative mechanism occasionally extends beyond potent neutralizing, narrowly reactive, class 1 antibodies (**Figs. 4C**, **4D** and **S6A**). We next examined the effects of anti-RBD antibodies at the polyclonal level using RBD-enriched and -depleted IgG fractions from six convalescent donors. Although both fractions bound to δ SARS-CoV-2 S proteins, only the RBD-enriched IgG antibodies enhanced Cv2F.14107 recognition (in 4 out of 6 donors; **Fig. 4E**). As expected, enhanced Cv2F.14107 binding to the spike did not occur for BA.2.86 and EG.5.1, likely due to the poor reactivity of anti-RBD antibodies against these variants (**Figs. S6B** and **S6C**). Principal component analysis demonstrated clear segregation between anti-RBD with narrow and broad reactivity based on antiviral functions (neutralizing, Fc-dependent effector and ACE2-mimetic activities) (**Fig. 4G**), suggesting that anti-RBD antibodies are typically either broadly reactive or potent polyfunctional immune effectors, but rarely both.

**Figure 4.**
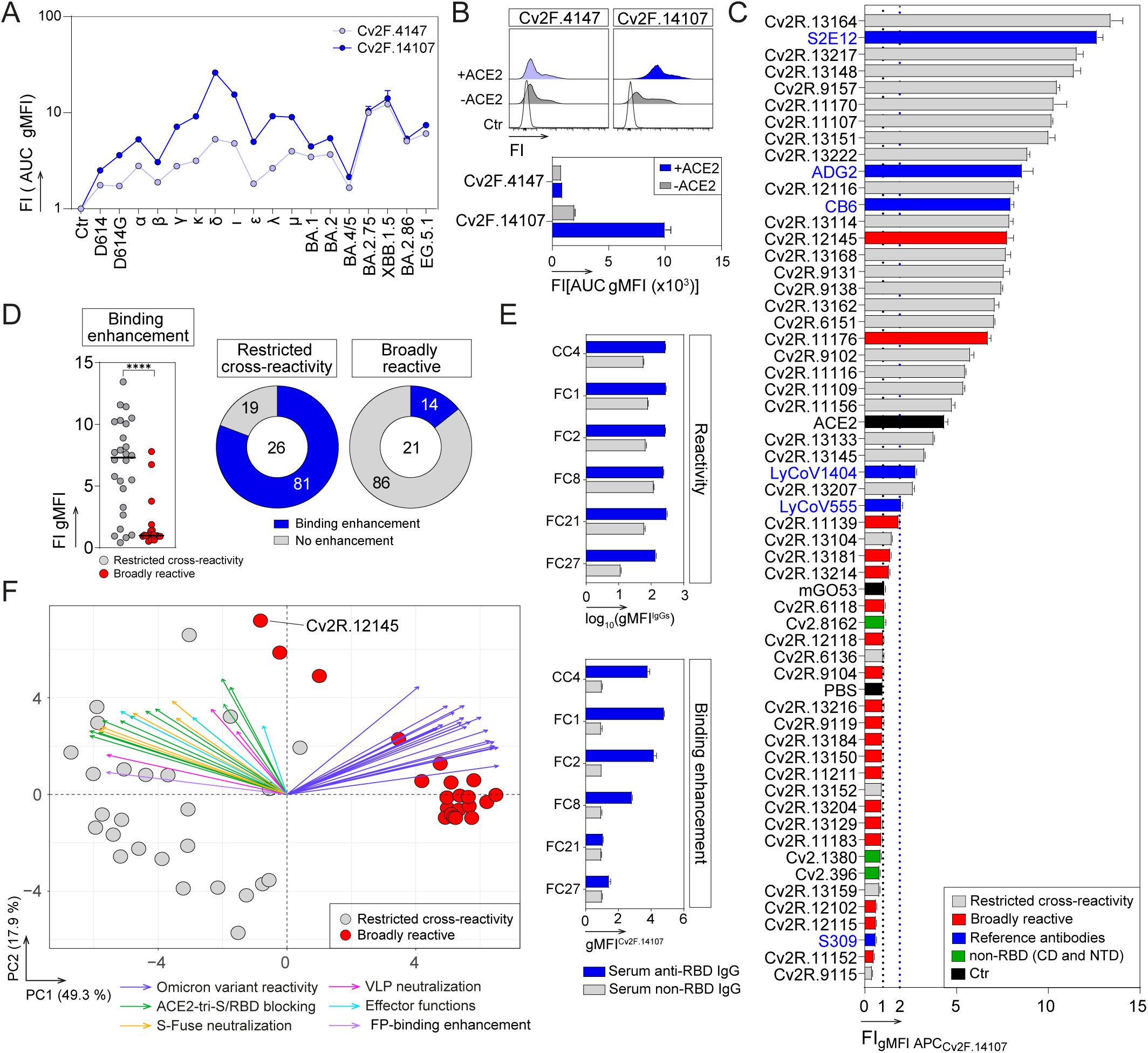
ACE2-mimetic activity of anti-RBD antibodies. (**A**) Graph comparing the reactivity of human anti-FP antibodies Cv2F.4147 and Cv2F.14107 against 293F-cell transfectants expressing the spike proteins from various SARS-CoV-2 variants. Ctr corresponds to the non-transfected cell control. Data represent means ± SD of duplicate values of the geometric mean fluorescence intensities (gMFI) fold increase compared to no antibodies. (**B**) Representative flow cytometry histograms (top) and bar graph (bottom) showing the flow cytometry binding of anti-FP antibodies to Delta spike-expressing 293F cells in the presence or absence of soluble recombinant ACE2 protein. Error bars in the graph represent the SD duplicate values for the fold increase (FI) of gMFI AUC compared to the no antibody condition. (**C**) Bar graph comparing the binding enhancement of anti-FP antibody Cv2F.14107 to Delta spike-expressing 293F cells in the presence of human anti-RBD antibodies. Data represent the mean ± SD of duplicate values. Dotted lines indicate the basal gMFI (black) and a two-fold increase (blue). Antibodies are ordered according to their ability to potentiate the binding of anti-FP antibody and are color-coded by group: broad- (red) or restricted- (gray) reactivity (defined in Fig. 2A), references (pink), non-RBD reactive (green), and controls (black). (**D**) Scatter plot (left) comparing the enhanced binding of Cv2F.14107 to spike measured in (C) according to the reactivity profiles of anti-RBD antibodies (broad, n=21 or restricted, n=26). Groups were compared using Mann-Whitney’s test. Donut charts (right) comparing the potency of anti-RBD antibodies to increase Cv2F.14107 binding (high >2; none or low ≤ 2 FI gMFI) according to their reactivity profiles. (**E**) Bar graphs comparing the Spike reactivity (left) and ACE2-mimetic capacity (right; enhanced Cv2F.14107 binding) measured by flow cytometry of total anti-RBD (blue) and non-RBD (grey) IgG antibodies purified from convalescent sera. Error bars represent the SD of duplicate values. (**F**) PCA 2D-plot showing the antiviral-related variables (arrows) discriminating human anti-RBD antibodies (n = 47). Dots represent single antibodies, color-coded based their reactivity profiles.

### Neutralizing activity of combined anti-RBD ACE2-mimetic and anti-FP antibodies

Cv2R.12145 is a rare, pan-reactive class 1 antibody that modestly neutralized pre-BA.2 SARS-2 variant (**Figs. 5A** and **5B**). Cv2R.12145 enhanced FP accessibility to Cv2F.14107 as efficiently as soluble ACE2 and previously described ACE2-mimetic antibody CB6^36,55^ (**Figs. 5C** and **S6D**). Unlike CB6, which lost its FP “unmasking” capacity against XBB.1.5, EG.5.1 and BA.2.86 due to altered reactivity, Cv2R.12145 still bound these variants and facilitated their recognition by Cv2F.14107 (**Figs. 5C** and **S7**). We next investigated whether combining Cv2R.12145 with Cv2F.14107 would result in additive or synergistic effects in neutralizing post-BA.1 variants. To this end, we conducted a synergy-scoring model analysis of *in vitro* neutralization data obtained on BA.5 and EG.5.1, using VLP reporter cell-based and S-Fuse assays, respectively. Antibodies in the cocktail showed no overall synergy, with total synergy δ-scores of 4.2 and 2.2, respectively; global synergy is typically defined by a δ-score of >10 (**Figs. 6D** and **6E**). However, δ-scores nearly doubled withing specific high concentration ranges for one or both antibodies (δ = 7.4 and δ = 4.25 for BA.5 and XBB.1.5, respectively) (**Figs. 6D** and 6**E**). These results indicate that, while Cv2R.12145 enhances the binding of Cv2F.14107 to SARS-CoV-2, their combined neutralization is primarily additive, with stronger localized effects at high concentrations.

**Figure 5.**
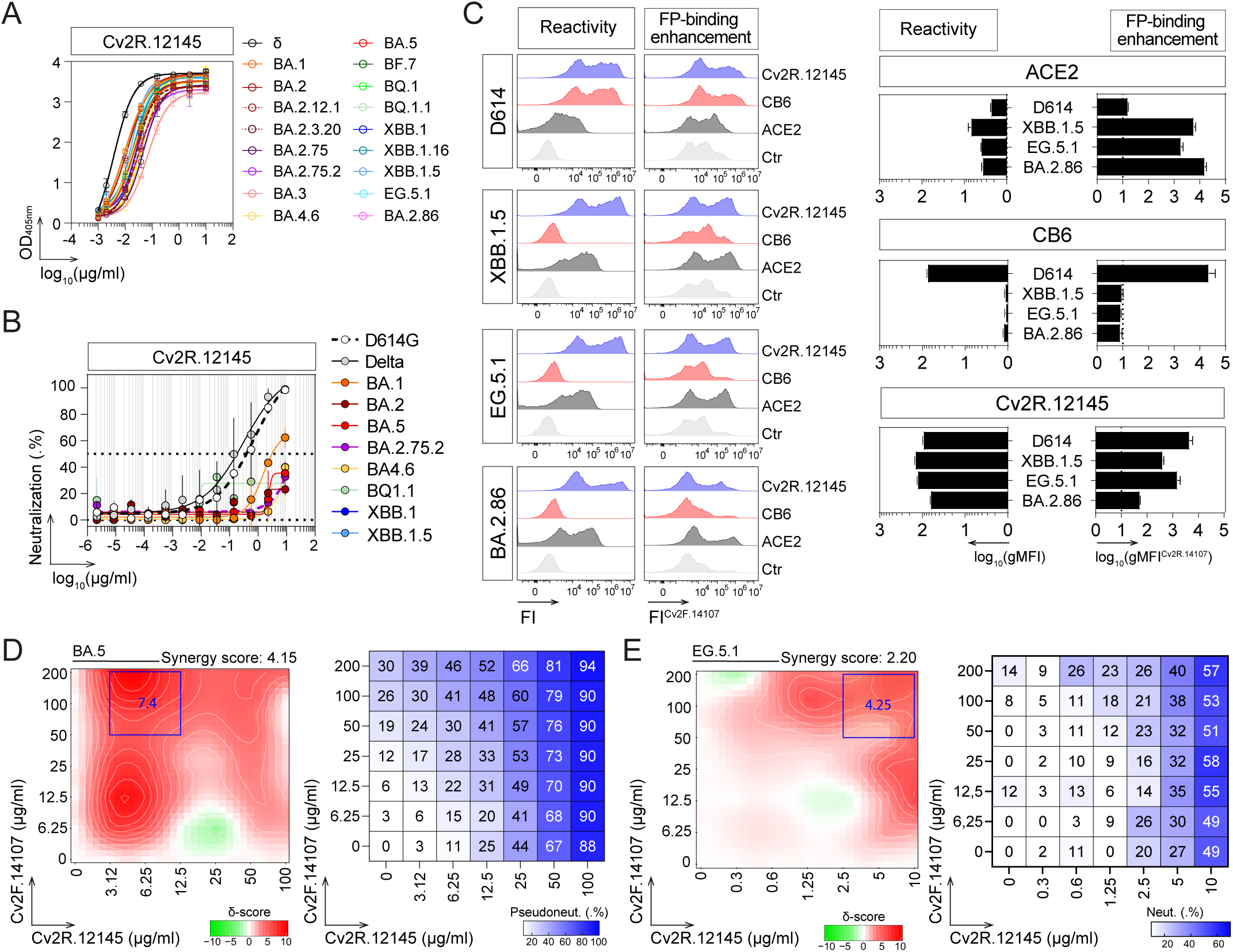
SARS-CoV-2 binding and neutralization by combined Cv2R.12145 and Cv2F.14107 antibodies. (**A**) Representative ELISA graph showing the binding curves of anti-RBD antibody Cv2R.12145 to purified RBD proteins from Delta and Omicron variants. Error bars represent the SD of duplicate values. (**B**) Graph showing the S-Fuse neutralization curves of Cv2R.12145 against SARS-CoV-2 D614G, Delta and Omicron variants. Error bars represent the SD of duplicate values from two intendent experiments. (**C**) Representative flow cytometry histograms (left) and bar graph (right) comparing the Delta spike binding and ACE2-mimetic capacity (right; enhanced Cv2F.14107 binding) of soluble recombinant ACE2 ectodomain, and anti-RBD antibodies Cv2R.12145 and CB6. Error bars in the graph represent the SD duplicate values. (**D**) Synergy maps (left) calculated on the dose-response neutralization matrices (heatmaps, right) of combined Cv2F.14107 and Cv2R.12145 against BA.5 using the VLP neutralization assay. Synergistic area are shown in red. Blue dotted square represent the highest synergy score (δ score). Global and highest local area synergy scores are indicated on each map. (**E**) Same as in (D) but for EG.5.1 viruses in the S-Fuse assay.

## DISCUSSION

While neutralization by anti-spike antibodies is the primary protective mechanism against COVID-19, their Fc-effector functions also contribute to immune protection, control, and clearance of infection^32,56^. Fc-effector functions of antibodies are governed by the interplay between paratope features (*i.e.*, epitope specificity and accessibility, binding affinity and stoichiometry, and overall binding geometry and topology) and Fc region properties (*i.e.*, isotype, subclass, glycosylation pattern)^57^. Fc-effector competent antibodies target multiple regions of the viral spike, including the RBD^32,56^. Here, we functionally profiled a panel of RBD-specific human monoclonal antibodies cloned from memory B cells of individuals infected with ancestral SARS-CoV-2. RBD-captured memory B cells produced both neutralizing and non-neutralizing antibodies. As expected from the escape and evolution^58,59^, most neutralizing antibodies lost their ability to bind and neutralize viruses in the Omicron lineage. Nonetheless, we identified a resilient class 1 anti-RBD neutralizing antibody, Cv2R.13162 - reminiscent of our previously reported broadly neutralizing antibody Cv2.3194^48,60^ - that maintains activity against Omicron variants up to XBB.1.5. Notably, class 1 neutralizing antibodies also mediated ADCC, ADCP, and CD against ancestral SARS-CoV-2 *in vitro*. Among these, Cv2R.12145 and Cv2.1176 stood out from the others by exhibiting broad reactivity against all RBD variants tested. However, most class 1 neutralizers likely fail to elicit Fc-effector functions against neutralization-resistant Omicron variants, as recently showed for infection- and vaccine-induced class 1/2 anti-RBD antibodies^61^. On the other hand, the majority of antibodies binding to non-neutralizing epitopes under lower selective pressure were cross-reactive against all tested RBD variants. Despite absent or weak neutralization and ADCP, these cross-reactive RBD-specific antibodies retained the capacity to induce CD and ADCC. Whether these anti-RBD antibodies would contribute to *in vivo* protection and viral clearance following infection with post-Omicron SARS-CoV-2 variants remains unclear. Notably, some vaccine-induced, non-neutralizing anti-RBD antibodies that mediate ADCC but not ADCP or CD activity have been shown to protect *in vivo* against lethal challenge with ancestral SARS-CoV-2^62^. More generally, in animal models, optimal *in vivo* prophylactic and/or therapeutic efficacy of neutralizing and non-neutralizing anti–SARS-CoV-2 antibodies, require intact Fc-mediated effector functions^28,62–65^. ACE2-mimetic anti-RBD neutralizing antibodies have previously shown to enhance anti-FP antibody reactivity and/or neutralization by inducing spike conformational changes^49–51^. We further show that contrary to broadly reactive SARS-CoV-2 antibodies binding outside the RBM, most class 1 neutralizers enhance antibody recognition of the FP on the spike protein. Combining the potent ACE2-mimetic Cv2R.12145 with the anti-FP Cv2F.14107 markedly increased Cv2F.14107 binding to spike and produced additive neutralization effects, but not the robust synergy previously reported for the anti-FP C77G12/anti-RBD S2E12 and anti-FP 76E1/anti-RBD CB6 combinations^49,50^. This is likely due to the low intrinsic potency of Cv2F.14107: despite targeting an epitope similar to 76E1, it exhibits substantially reduced binding affinity and neutralizing activity against SARS-CoV-2.

Neutralizing antibodies impose strong selective pressure for viral escape, driving SARS-CoV-2 evolution that culminated in the emergence of highly mutated Omicron lineages^58^. As a result, potent anti-RBD neutralizers - particularly class 1 antibodies - have largely lost their neutralization capacity against Omicron variants^66–69^. Our study suggests that viral evolution did not only confer resistance to neutralization by class 1 antibodies, but also more broadly compromised Fc effector and allosteric antiviral functions. Loss of these latter activities may further reduce the protective efficacy of SARS-CoV-2 antibodies. Disruption of Fc-mediated effector functions of anti-RBD neutralizing antibodies has been shown to markedly impair their prophylactic and therapeutic efficacy in rodent models^56,63^. Thus, SARS-CoV-2 can escape potent, polyfunctional neutralizers. In contrast, we show that anti-RBD antibodies targeting epitopes outside the RBM, with broad recognition across Omicron lineages, can still promote ADCC and complement deposition. This may explain why anti-RBD antibody effector activities measured serologically in convalescents exposed to ancestral SARS-CoV-2 are not completely abrogated against pre-Omicron variants^70^. Effector-competent non-neutralizing antibodies, including those targeting the RBD, can efficiently protect mice against SARS-CoV-2 infection^62,65^. It is generally accepted that Fc mediated antibody effector functions play an important role once infection is established, but have a more limited impact on sterilizing immunity than neutralization^31,32,56^. Whether cross-reactive, non-neutralizing anti-RBD effector responses elicited by ancestral infection contribute substantively to protection against Omicron-mediated COVID-19 remains unclear. Of note, Wuhan-based vaccines that cross-protect against severe COVID-19 elicit anti-spike antibodies with durable but variable Fc-effector activities^71^, which appear to be largely preserved against Omicron lineages, albeit to varying degrees depending on the specific variant^34,72,73^.

### Limitations of the study

We employed *in vitro* surrogate assays to evaluate the Fc-dependent effector functions of human anti-RBD antibodies using ancestral SARS-CoV-2 proteins. Our data would benefit from additional analyses using primary infected target and effector cells, inclusion of further viral variants (*e.g.*, Omicron strains) and anti-RBD antibody classes, as well as *in vivo* confirmation in relevant animal models. Assessing the *in vivo* Fc-mediated antiviral activities of anti-RBD antibodies - including engineered variants with altered Fcγ receptor binding - may provide valuable insights into the contribution of Fc-dependent effector functions to infection protection and resolution. Although most anti-RBD monoclonals were produced as native IgG1, we did not generate recombinant versions of non-IgG1 antibodies (*e.g.*, IgG3 and IgA1), precluding assessment of Fc-dependent effector functions in their original subclass. In addition, the glycosylation profiles of these recombinant antibodies may differ from those of naturally produced IgGs, potentially modulating their Fc-effector activities.

## RESOURCE AVAILABILITY

### Lead Contact

Requests for resources and reagents should be directed to and will be fulfilled by the lead contact, Hugo Mouquet.

### Materials Availability

Request for reagents will be made available by the lead contact with a Material Transfer Agreement.

### Data and code Availability

This paper does not report original code. Data used for the analysis is available in supplemental tables. Any additional information required to reanalyze the data reported in this paper is available from the lead contact upon reasonable request.

## ACKNOWLEDGMENTS

We are grateful to all convalescent donors who consented to participate in this study, as well as to everyone who contributed to the CORSER and French COVID studies. We thank Sandrine Schmutz and Sophie Novault for their help with single-cell sorting (CB-UTechS, Institut Pasteur), and Dr Lubka Roumenina (Centre de Recherche des Cordeliers, France) for providing pre-pandemic sera for complement-related experiments. The French COVID Cohort, sponsored by the INSERM, was funded by the REACTing consortium, the Ministry of Health and Social Affairs, the Ministry of Higher Education and Research dedicated COVID-19 fund and Programme Hospitalier de Recherche Clinique (no. 20-0424). This work was supported by grants from the Agence Nationale de la Recherche REACTing COVID-19 (#20RR028-00), the European Commission Horizon 2020 program (RECoVER project, #101003589), the Institut Pasteur Task Force COVID-19 (2019-NCOV THERAMAB project), and the Fondation de France (#00106077). P.R was supported by a two-years Pasteur-Roux-Cantarini fellowship (Institut Pasteur).

## AUTHOR CONTRIBUTIONS

H. M. conceived and supervised the study. P. R., C. P., T. B., M. B., W.H. B., I. S., F. G., and O. S., H. M. performed and/or analyzed the experiments. French COVID and CORSER Cohort Study Groups provided human samples. P. R. and H.M wrote the manuscript with contributions from all the authors.

## SUPPLEMENTAL INFORMATION

**Figure S1.**
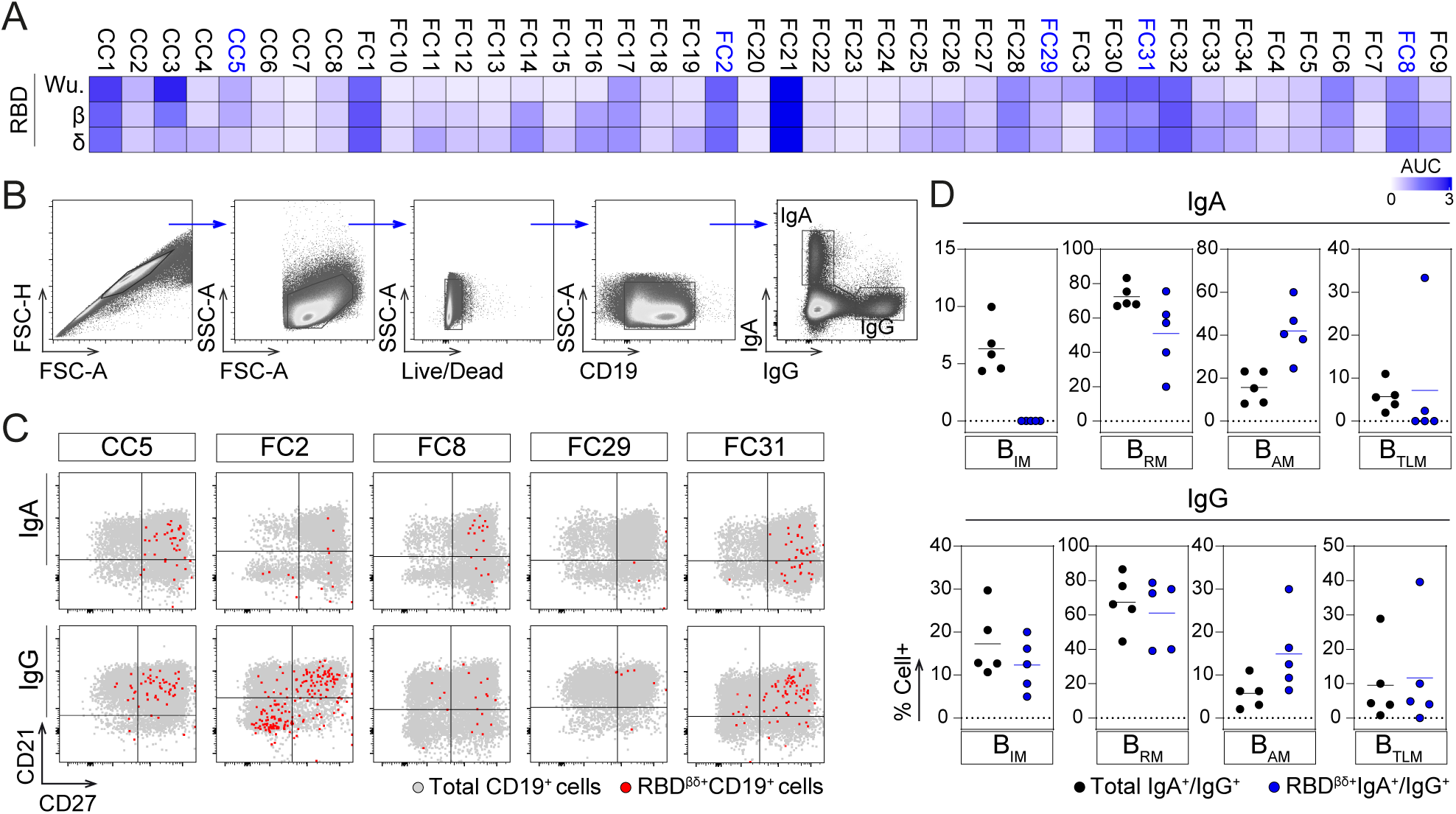
RBD-seroreactivity and -memory B-cell sorting. (**A**) Heatmap comparing the ELISA IgG seroreactivity of COVID-19 convalescents against purified recombinant Wuhan (Wu.), Beta (β), and Delta (δ) RBD proteins. Cells are color-coded according to the area under curve (AUC) values obtained by ELISA titrations. Blue-labeled donor codes indicate the samples selected for the flow cytometry B-cell sorting. (**B**) Flow cytometry plots presenting the gating strategy used to identify and single-cell sort SARS-CoV-2 RBDβδ^+^-binding memory B cells. SSC, side scatter; FSC, forward scatter. (**C**) Flow cytometric plots showing for each donor the overlays of CD21 and CD27 stainings on total CD19^+^ cells (gray) with SARS-CoV-2 RBDβδ^+^ cells (red). (**D**) Scatter plots comparing the distribution of B-cell subsets between total (black) and SARS-CoV-2 RBDβδ^+^ (blue) IgG^+^ and IgA^+^ memory B cells (n = 5 donors): B_IM_ (intermediate memory, CD27^−^CD21^+^), B_RM_ (resting memory, CD27^+^CD21^+^), B_AM_ (activated memory, CD27^+^CD21^−^), and B_TLM_ (tissue-like memory, CD27^−^CD21^−^).

**Figure S2.**
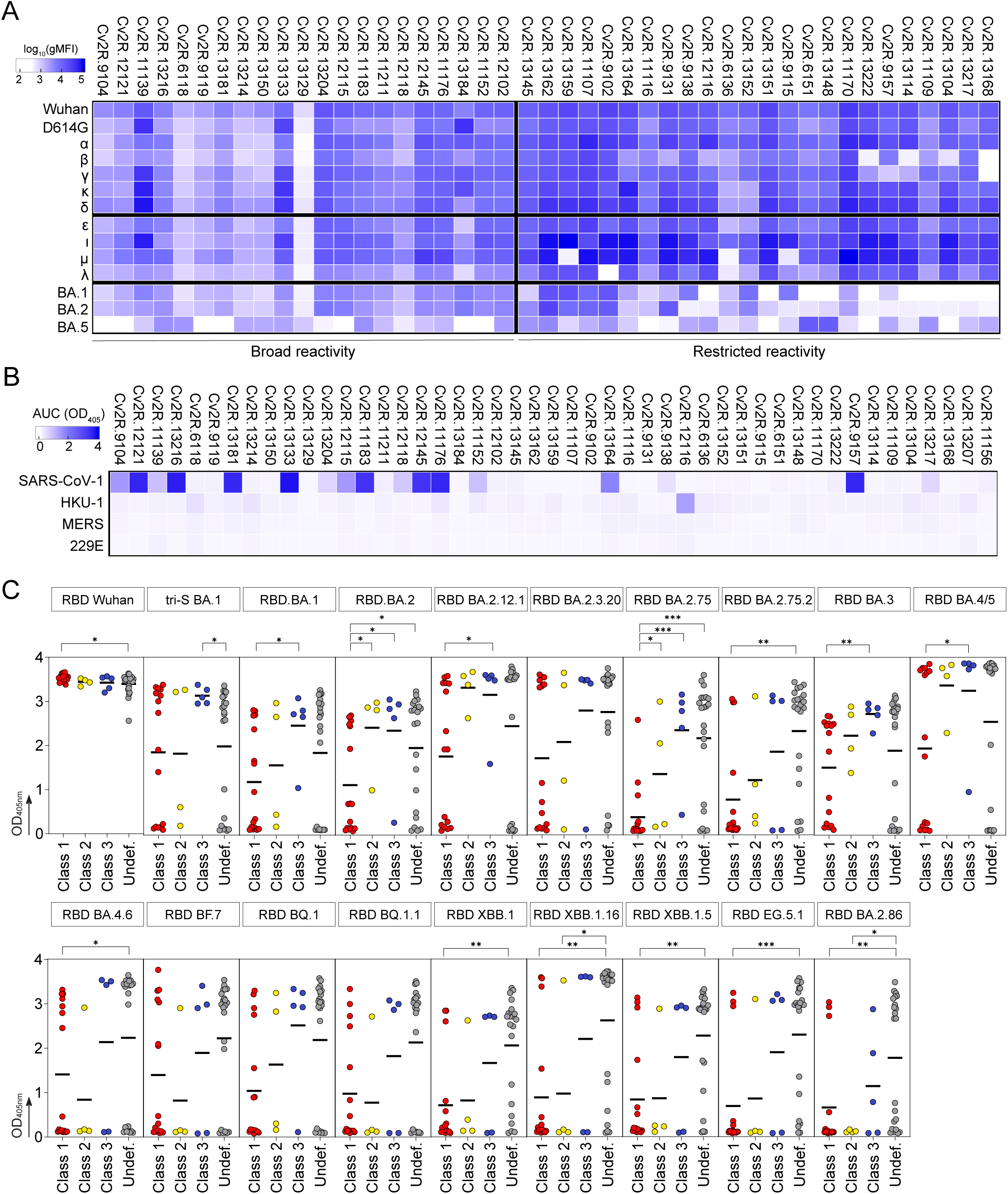
Reactivity profiling of human anti-RBD antibodies. (**A**) Heatmap comparing the reactivity profiles of anti-RBD across on 293F-cell transfectants expressing spike proteins from various SARS-CoV-2 variants. Cells are color-coded according to log_10_ geometric mean fluorescence intensity (gMFI) values, with dark blue colors indicating strong reactivities. Antibodies were categorized as either broadly reactive (red) or with a more restricted cross-reactivity (grey). (**B**) Heatmap comparing the ELISA reactivity of anti-RBD antibodies - measured as AUC from antibody titrations - against purified recombinant SARS-CoV-1, HKU-1, MERS-CoV, 229E tri-S proteins. Dark blue cells indicate high reactivities. Data correspond to mean of duplicate values. (**C**) Scatter plots comparing the anti-RBD antibody reactivities against RBD proteins from Wuhan and Omicron variants determined by ELISA. Antibody groups were defined based on RBD classes [1, 2, 3 and undefined (undef.)] and compared using Mann-Whitney’s test. ***, p < 0.001; **, p < 0.01; *, p < 0.05.

**Figure S3.**
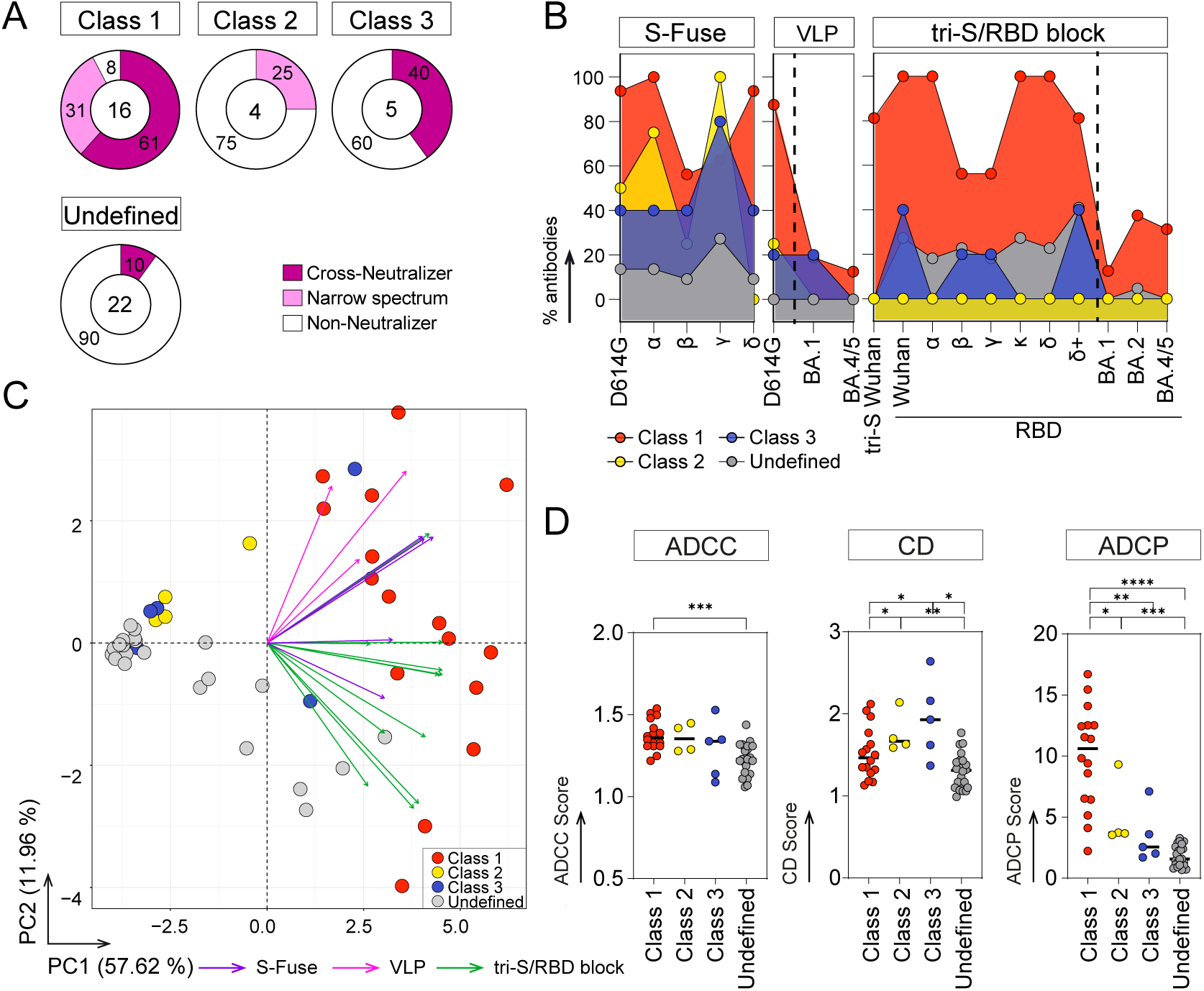
Functional profiling of anti-RBD antibody classes. (**A**) Donut charts comparing the distribution of cross-neutralizing (purple), narrow spectrum (pink), or non-neutralizing (white) antibodies according to their RBD class (1, 2, 3 and undefined). (**B**) Graph comparing the neutralization profiles of anti-RBD antibodies as determined in the S-fuse and VLP neutralization assays, and ACE2-tri-S or -RBD blocking ELISA assay according to their RBD class as defined in Fig.2B. (**C**) PCA 2D-plot showing the neutralization variables (arrows; S-fuse, VLP, ACE2-tri-S or -RBD blocking assays) discriminating human anti-RBD antibodies (n = 47). Dots represent single antibodies, color-coded based on their RBD class. (**D**) Scatter plots comparing the ADCC, CD, and ADCP activities of anti-RBD specific antibodies according to their RBD class. Groups were compared using Mann-Whitney’s test. ****, p < 0.0001; ***, p < 0.001; **, p < 0.01; *, p < 0.05.

**Figure S4.**
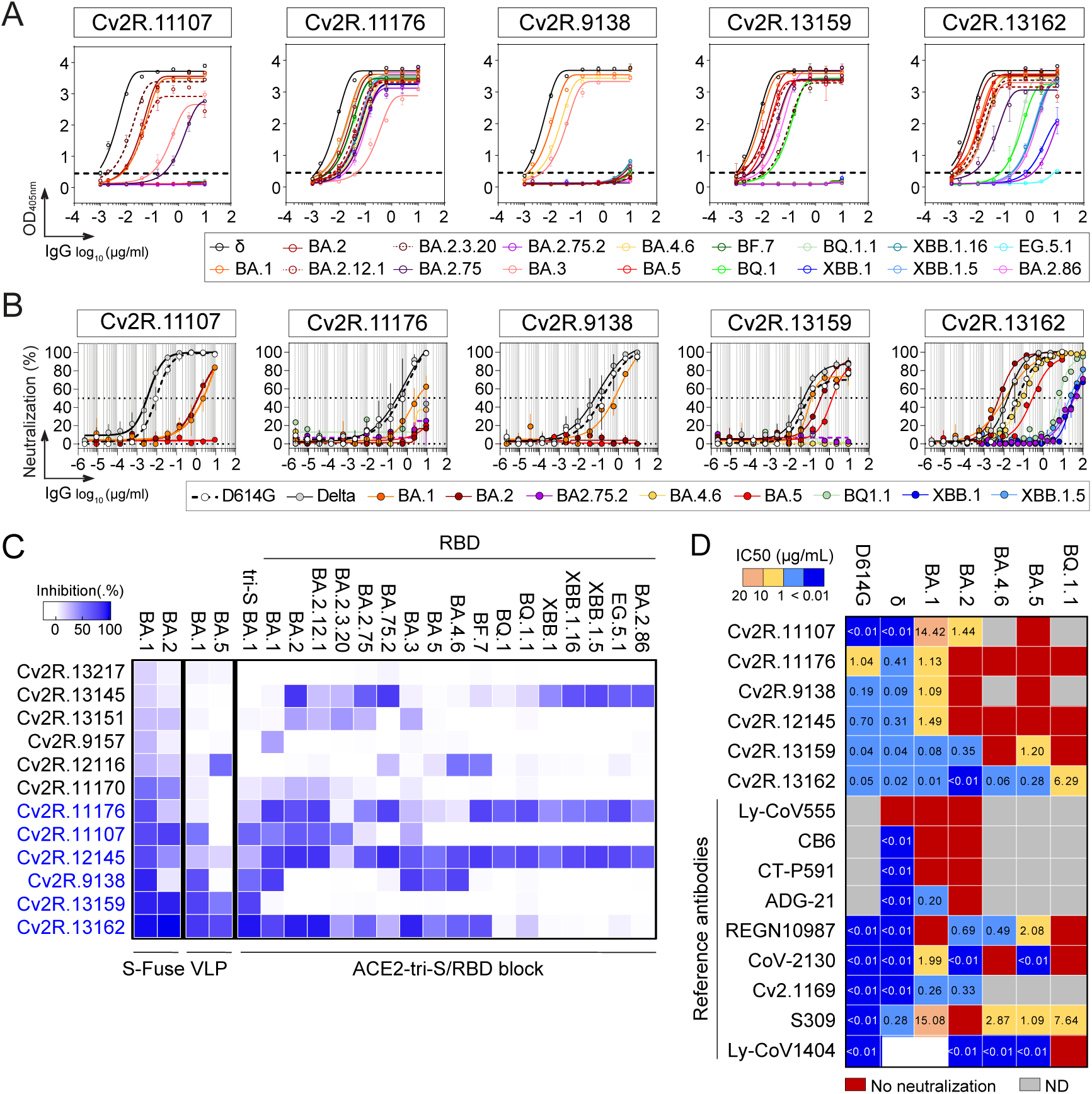
SARS-CoV-2 cross-reactivity and -neutralization profiles of anti-RBD neutralizers. (**A**) ELISA graph comparing the binding curves of the five selected potent anti-RBD IgG antibodies against SARS-CoV-2 RBD proteins from Delta and Omicron variants Error bars represent the SD of duplicate values. (**B**) Same as in (A) but for in vitro neutralization in the S-Fuse assays using authentic viruses. (**C**) Heatmap comparing the neutralizing activities of potent anti-RBD antibodies as determined in the S-fuse and VLP neutralization assays, and ACE2-tri-S or -RBD blocking ELISA assay. Mean of duplicate values are shown. Dark blue colors indicate strongest inhibition percentages, while white corresponds to no inhibition. Blue-labeled antibodies (>70% SARS-CoV-2 BA.1 neutraliza-tion) were considered as the most potent. (**D**) Table comparing the IC_50_ values of the 6 most potent cross-neutralizing anti-RBD antibodies and reference antibodies against SARS-CoV-2 D614G, Delta, and Omicron variants measured in the S-Fuse neutralization assay as shown in (A). ND, not determined.

**Figure S5.**
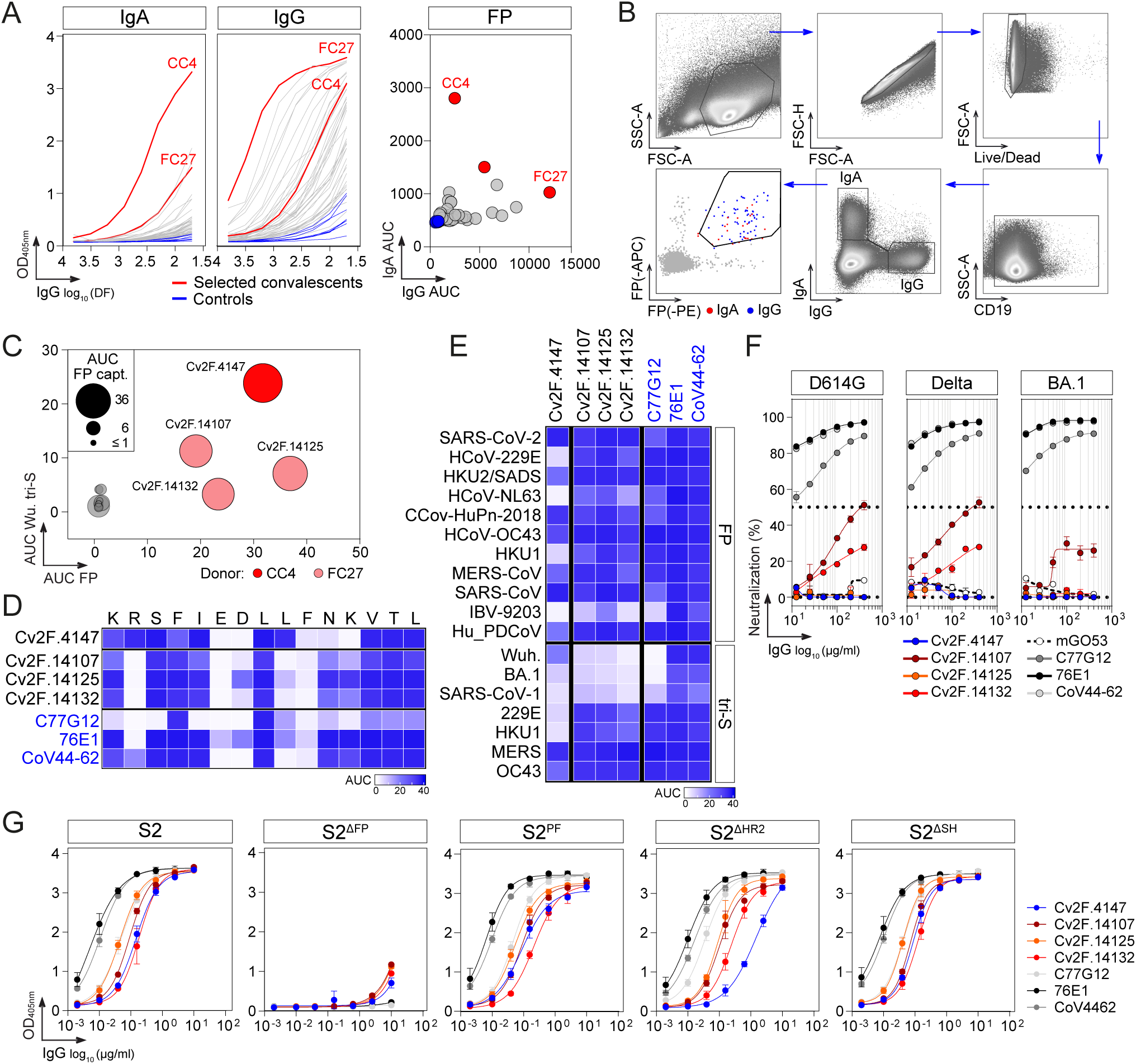
Human SARS-CoV-2 FP-specific memory B-cell antibody cloning and characterization. (**A**) ELISA graphs (left) comparing the IgG and IgA seroreactivities from COVID-19 convalescent indivi-duals (n=42, non-blue lines) against SARS-CoV-2 fusion peptide (FP). Mean of duplicate values are shown. Red and blue lines correspond to the convalescent donors selected for single B-cell sorting (CC4 and FC27) and healthy individual controls (n=8), respectively. DF, dilution factors. Dot plot (right) compa-ring IgG and IgA FP-seroreactivity, shown as AUC values derived from the ELISA titrations. (**B**) Flow cytometry plots presenting the gating strategy used to identify and single-cell FP-binding memory B cells from the PBMC of CC4 and FC27 donors. (**C**) Bubble plot comparing the ELISA binding of 17 human antibodies cloned from SARS-CoV-2 FP-captured IgG^+^ and IgA^+^ memory B cells of CC4 (blue) and FC27 (red) donors. The y-axis shows binding to Wuhan tri-S protein; the x-axis and bubble size represent binding to FP peptide measured by classical indirect and capture ELISA (capt.), respectively. Grey bubbles indicate non-reactive antibodies from both donors. Values correspond to mean AUC values calculated from dupli-cate titrations. (**D**) Heatmap comparing the ELISA binding of anti-FP peptides and reference antibodies to alanine-mutated FP peptides (mutated residues at the top). Values correspond to mean AUC values calcu-lated from duplicate titrations, with dark blue cells indicating strong binding. (**E**) Same as in (D) but for fusion peptides and tri-S proteins from different coronaviruses. (**F**) Graphs comparing the SARS-CoV-2 neutralizing activity of anti-FP and reference antibodies as measured in the S-Fuse neutralization assay. Error bars represent the SD of duplicate values. (**G**) ELISA graphs comparing the binding of anti-FP and reference antibodies to pre- and post-fusion trimeric SARS-CoV-2 S2 subunits (S2 and S2^PF^), and S2 protein versions deleted from the fusion peptide (S2^ΔFP^), heptad-repeat 2 (S2^ΔHR2^) or stem-helix (S2^ΔSH^). Error bars correspond the SD of duplicate values.

**Figure S6.**
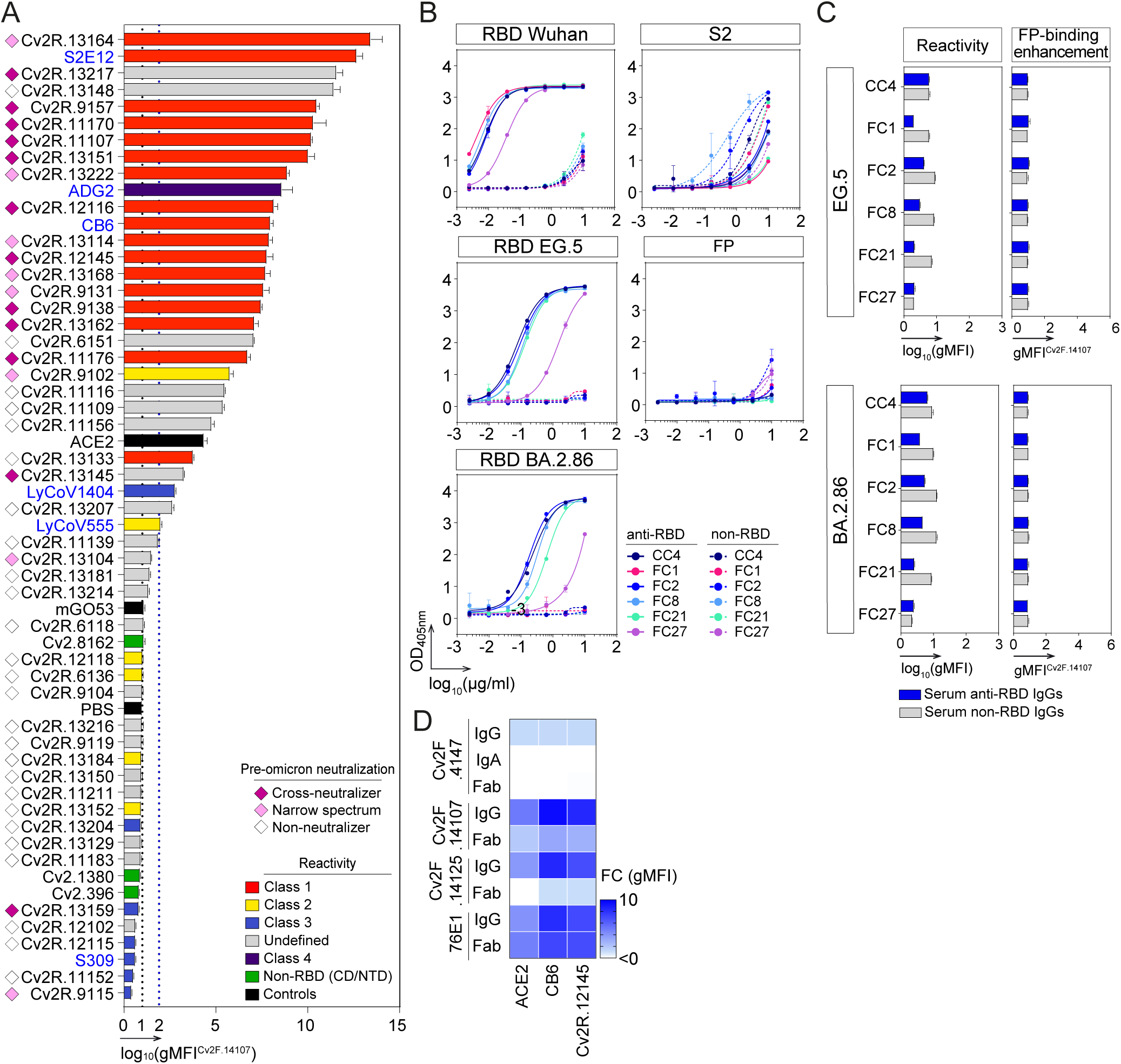
ACE2-mimetic activity of anti-RBD antibodies. (**A**) Bar graph comparing the enhanced binding of anti-FP Cv2F.14107 to Delta tri-S-expressing 293F cell transfectants in the presence of anti-RBD antibo-dies. Bars are color-coded according to the RBD class and show the means ± SD of duplicate values. Dotted lines indicate the basal gMFI (black) and a two-fold increase (blue). Diamond symbols on the left are color-coded according to the neutralization profiles on pre-Omicron viruses (see Fig.3A): cross-neutralizing (purple), narrow spectrum (pink), or non-neutralizing (white). (**B**) ELISA graphs comparing the binding of total anti-RBD and non-anti-RBD IgG antibodies purified from six convalescent sera against Wuhan, EG.5 and BA.2.86 RBD proteins, Wuhan S2 subunit and FP peptide. Error bars correspond to the SD of duplicate values. (**C**) Bar graphs comparing the EG.5 and BA2.86 Spike binding (left) and ACE2-mimetic capacity (right; enhanced Cv2F.14107 binding) measured by flow cytometry of total anti-RBD (blue) and non-RBD (grey) IgG antibodies purified from convalescent sera. Error bars represent the SD of duplicate values. (**D**) Heatmap comparing the ACE2-mimetic capacity of soluble ACE2, CB6, and Cv2R.12145 measured by flow cytometry on Delta spike-expressing 293F transfectants as enhanced binding of anti-FP antibodies (Cv2F.4147, Cv2F.14107, CV2F.14125 and 76E1) produced as recombinant IgG, IgA or Fab. Cells are color-coded based on the mean fold change (FC) of duplicate gMFI values relative to the no antibody control (means of duplicate values). Darker blue indicates a higher fold change signal intensity

**Figure S7.**
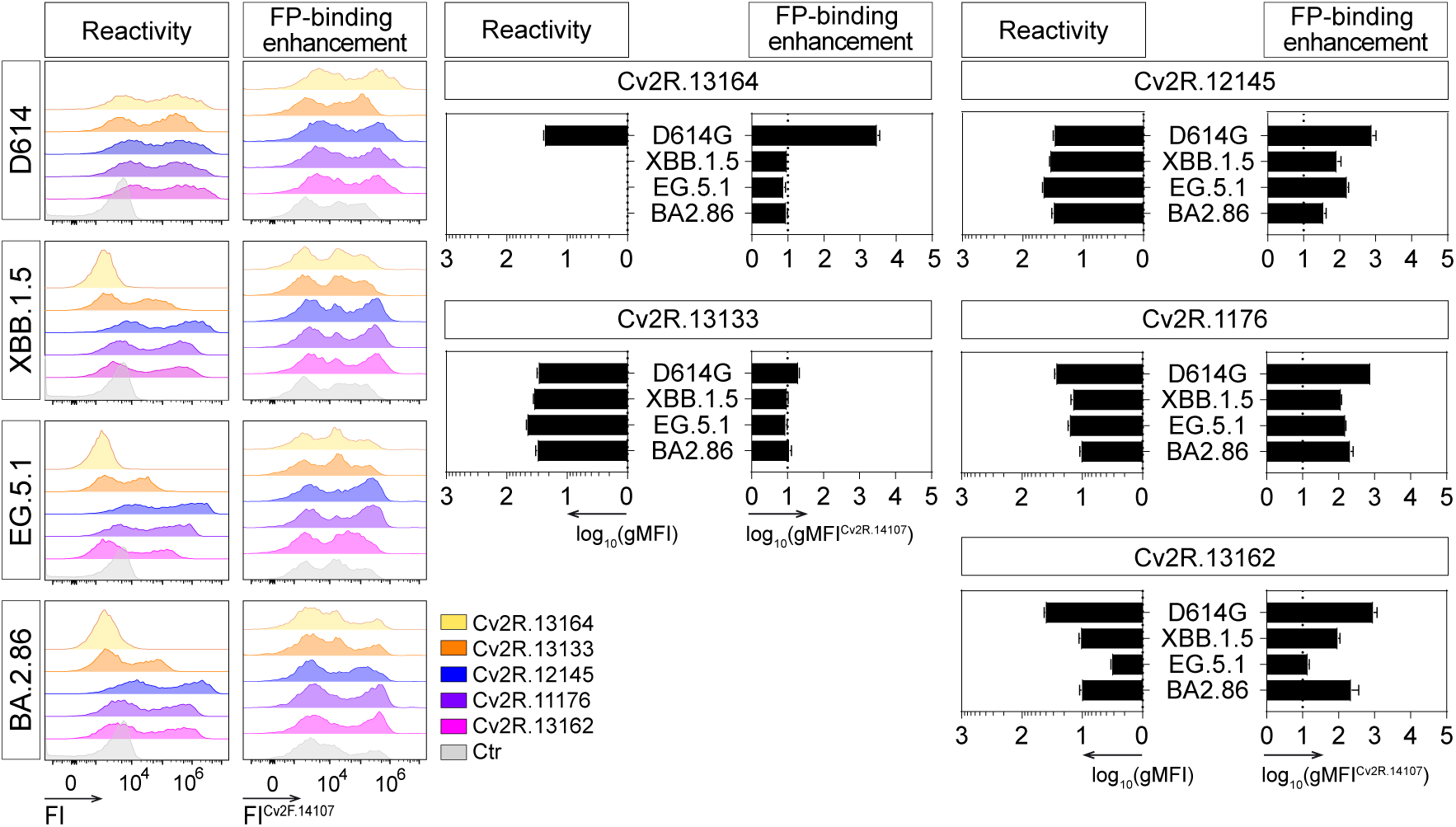
ACE2-mimetic activity of broadly reactive anti-RBD antibodies. Represen-tative flow cytometry histograms (left) and bar graph (right) comparing the spike binding and ACE2-mimetic capacity (based on enhanced Cv2F.14107 binding) of 5 broadly reactive anti-RBD IgG antibodies as determined by flow cytometry using 4 different SARS-CoV-2 strains. Error bars in graphs represent the SD of duplicate values. Ctr, no antibody control.

**Table S1.**
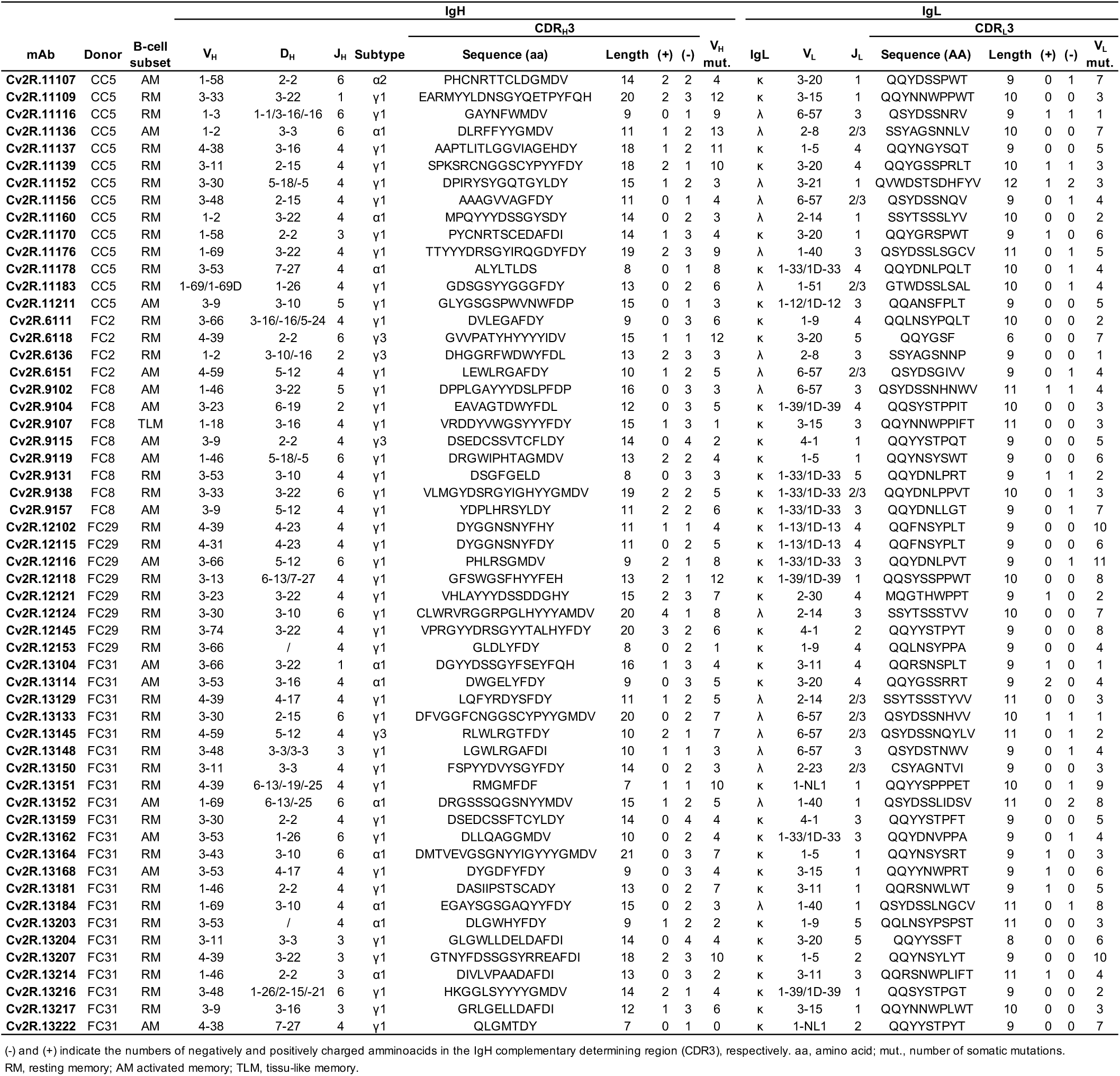
Immunoglobulin gene repertoire of human SARS-CoV-2 RBD memory B-cell antibodies.

**Table S2.**
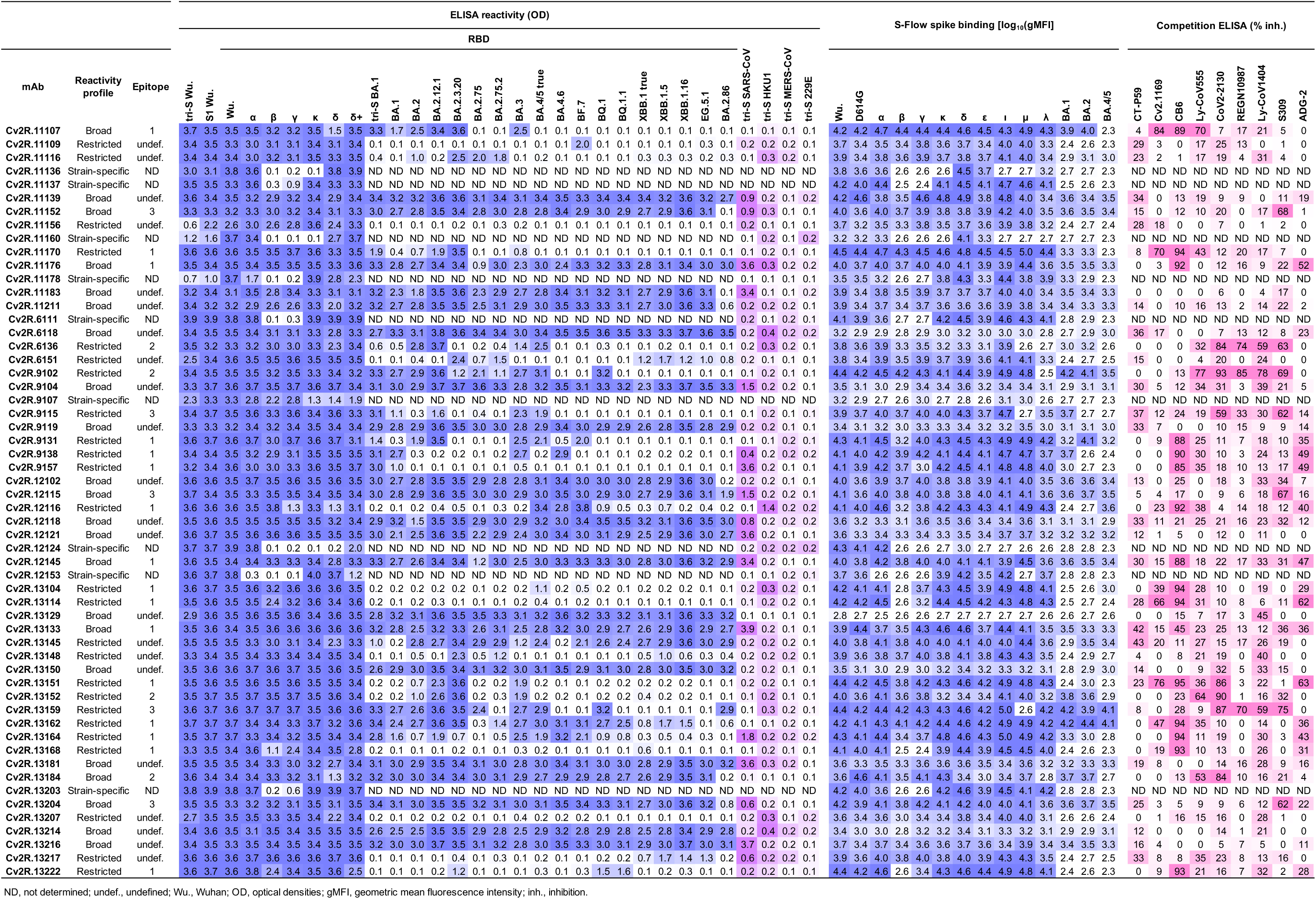
Binding profiles of human SARS-CoV-2 RBD memory B-cell antibodies.

**Table S3.**
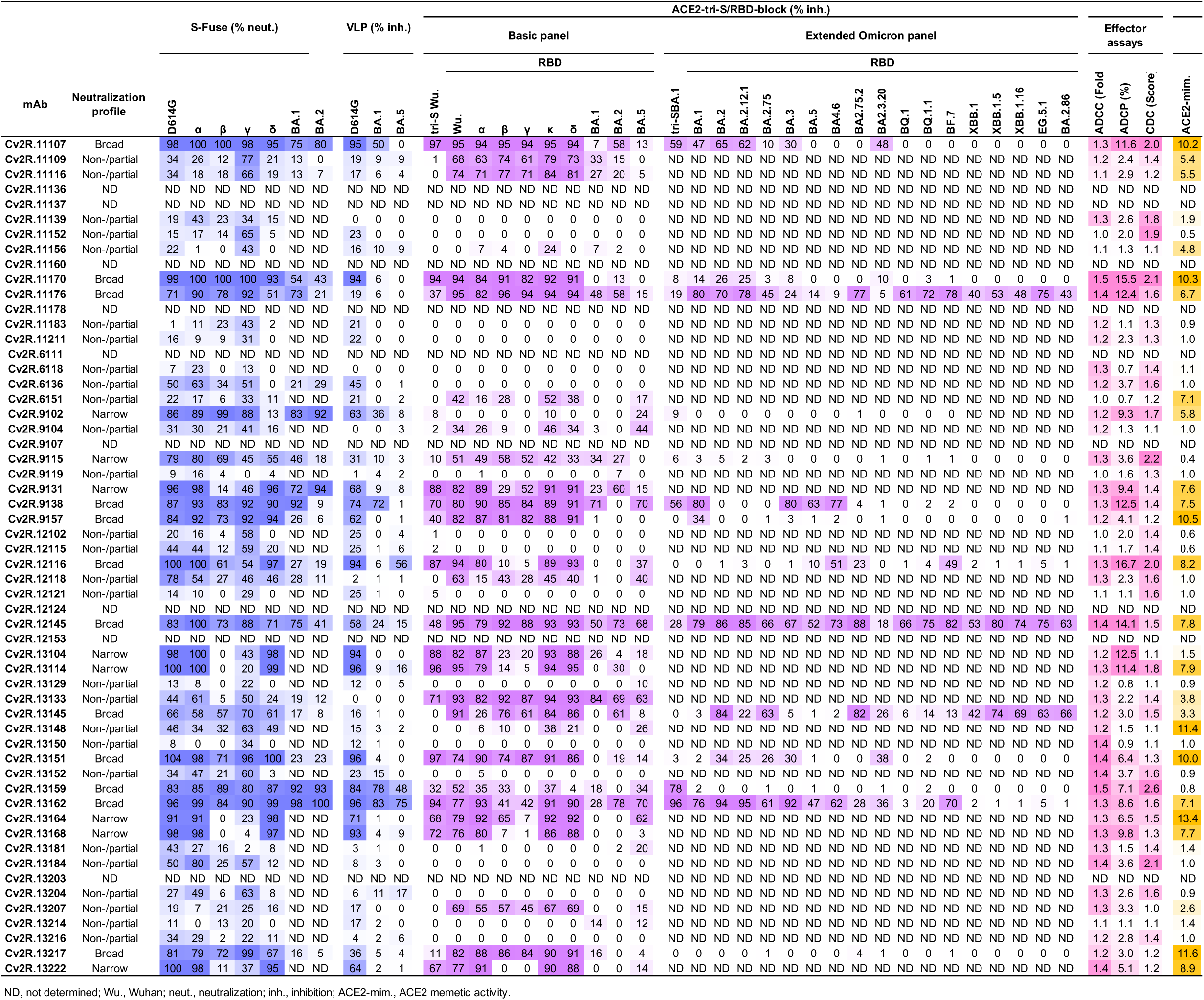
Antiviral functions of human SARS-CoV-2 RBD memory B-cell antibodies.

## STAR★METHODS

### EXPERIMENTAL MODEL AND SUBJECT DETAILS

#### Human samples

Biobanked blood-derived samples from Wuhan COVID-19 convalescent donors (n=42) were previously obtained from the CORSER and French COVID cohorts^38^. These samples were collected from participants infected during the first the first epidemic wave in France, with blood sampling conducted in April 2020. Both cohorts were established in accordance with French legislation and regulations and following ethical approval from the relevant authorities. The CORSER study (NCT04325646) received ethical approval from the Comité de Protection des Personnes Ile-de-France III. The French COVID study was approved by the regional institutional review board (Comité de Protection des Personnes Ile-de-France VII, Paris, France) and conducted in accordance with European guidelines and the Declaration of Helsinki. All participants provided written informed consent to participate in this study, and data were collected in a pseudo-anonymized form using subject codes.

### METHOD DETAILS

#### Viruses

Titrated SARS-CoV-2 viral stocks for the following strains used in the *in vitro* S-Fuse neutralization assay were prepared in the Viral and Immunity Unit (Institut Pasteur) as previously described^69,74–78^: D614G (hCoV-19/France/GE1973/2020; GISAID ID: EPI_ISL_414631) BetaCoV/France/IDF0372/2020 (GISAID ID: EPI_ISL_406596) (National Reference Centre for Respiratory Viruses, Institut Pasteur), α (GISAID ID: EPI_ ISL_735391), β (B.1.351; GISAID ID: EPI_ISL_964916), γ variant (P.1.; hCoV-19/Japan/TY7-501/2021; GISAID ID: EPI_ISL_833366)^79^ δ (B.1.617.2; GISAID ID: EPI_ISL_2029113), BA.1 (GISAID ID: EPI_ISL_6794907), BA.2 (GISAID ID: EPI_ISL_10654979), BA.5 (GISAID ID: EPI_ISL_13660702), BA.2.75.2 (GISAID ID: EPI_ISL_15731524), BA.4.6 (GISAID ID: EPI_ISL_15729633, BQ.1.1 (GISAID ID: EPI_ISL_15731523, XBB.1 (GISAID ID: EPI_ISL_15619797), XBB.1.5 (GISAID ID: EPI_ISL_1635849) and EG.5.1 (GISAID ID: EPI_ISL_17949406).

#### Viral proteins and peptides

Codon-optimized nucleotide fragments encoding SARS-CoV-2 S2 subunit deleted from the FP, HR2, or SH domain, followed by a Foldon trimerization motif and C-terminal tags (8xHis-tag, Strep-tag, and AviTag) were synthesized and cloned into pcDNA3.1/Zeo(+) expression vector (Thermo Fisher Scientific). Already cloned pcDNA3.1/Zeo(+) vectors coding for the following tagged proteins were used to produce recombinant proteins by transient transfection of exponentially growing Freestyle 293-F suspension cells (Thermo Fisher Scientific) as previously described^80^: SARS-CoV-2, SARS-CoV-1, MERS-CoV, HKU1-CoV, 229E-CoV, SARS-CoV-2 BA.1 spike (HexaPro; S_6P; S) ectodomains; SARS-CoV-2 S2 subunit; tagged Wuhan, BA.1, BA.2, BA.2.12.1, BA2.75, BA2.75.2, BA.3, BA4/5, BA4.6, BF.7, BQ.1, BQ.1.1, XBB.1, XBB.1.16, XBB.1.5, and BA.2.86 RBD proteins. Proteins were purified from culture supernatants by high performance chromatography using the Ni Sepharose Excel Resin according to the manufacturer’s instructions (GE Healthcare), dialyzed against PBS using Slide-A-Lyzer dialysis cassettes (Thermo Fisher Scientific), quantified using NanoDrop 2000 instrument (Thermo Fisher Scientific). AviTagged tri-S and RBD proteins were biotinylated using the Enzymatic Protein Biotinylation Kit (Sigma-Aldrich) following the manufacturer’s instructions. For single B-cell FACS sorting, biotinylated RBD-β and RBD-δ proteins were coupled with respectively PE- and APC-conjugated streptavidin (Thermo Fisher Scientific), respectively; biotinylated fusion peptide (FP-biot; KRSFIEDLLFNKVTLADAGFIK, GenScript Biotech) was coupled to both. The library of 15-mer alanine-mutated SARS-CoV-2 FP (original reference: KRSFIEDLLFNKVTL, n=15), and FP from SARS-CoV-2 (PSKRSFIEDLLFNKVTLADA), SARS-CoV-1 (PTKRSFIEDLLFNKVTLADA), HCoV-229E (VAGRSAIEDILFSKIVTSGL), MERS-CoV (RSARSAIEDLLFDKVTIADP), HCoV-HKU1 (SSRSLLEDLLFNKVKLSDV), SADS-CoV (FESRSVIEDLLFSKIETTGP), HCoV-NL63 (IAGRSALEDLLFSKVVTSGL), HCoV-OC43 (ASSRSAIEDLLFDKVKLSDV), CCov-HuPn-2018 (RKYRSAIEDLLFDKVVTSGL), IBV-9203 (PRRRSFIEDLLFTSVESVGL) and Hu-PDCoV (LGGRSAIEDLLFNKVVTSGL) were synthetized and desalted (GenScript Biotech).

#### Reference and control antibodies

Purified parental research IgG1 versions of benchmarked anti-RBD antibodies (REGN10987^40^, CB6^41^, LY-CoV555^42^, CT-P59^43^, COV2-2130^44^, ADG-2^45^, Ly-CoV1404^13^ and S309^46^), anti-FP antibodies (C77G12^37^, 76E1^36^, CoV44-62^54^) were produced as described below following cloning of synthetic DNA fragments encoding immunoglobulin variable domains (Thermo Fisher Scientific). Non-SARS-CoV-2 antibody mGO53^81^ (non-polyreactive) was used as a negative control.

#### Single B-cell FACS sorting and expression-cloning of antibodies

Human B cells were isolated from biobanked PBMCs using human CD19 MACS following manufacturer’s procedures (Miltenyi Biotec). B cells were simultaneously incubated for 30 min at 4°C with an extemporaneously prepared mix of PE-coupled RBD-β and APC-coupled RBD-δ proteins (2 µg each) or with PE- and APC-coupled FP (2 µg each). Cells were washed once with 1% FBS-PBS (FACS buffer), stained using LIVE/DEAD aqua fixable dead cell stain kit (Molecular Probes; Thermo Fisher Scientific) and incubated 30 min with a cocktail of mouse anti-human antibodies: CD19 Alexa 700 (HIB19; BD Biosciences), CD21 BV711 (or BV421; B-ly4; BD Biosciences), CD27 PE-CF594 (M-T271; BD Biosciences), IgG BV786 (G18-145; BD Biosciences), IgA FITC (IS11-8E10; Miltenyi Biotec), and β7 integrin BV421 (or BUV395; FIB504, BD Biosciences). Cells were then washed and resuspended in 1 mM EDTA FACS buffer. Single alive RBDβ/δ^+^CD19^+^IgA^+^ or IgG^+^ and PE/APC-FP^+^CD19^+^IgG^+^ cells were sorted into 96-well PCR plates using either a FACSAria Fusion or a FACSymphony™ S6 cell sorter (Beckton Dickenson). Single-cell cDNA synthesis using SuperScript IV reverse transcriptase (Thermo Fisher Scientific) followed by nested-PCR amplifications of IgH, Igκ, and Igλ genes were performed as previously described^82,83^. Purified digested PCR products were cloned into human Igγ1-, Igκ-, or Igλ-expressing vectors (GenBank# LT615368.1, LT615369.1, and LT615370.1, respectively), as previously described^83^. Recombinant antibodies were produced by transient co-transfection of Freestyle 293-F suspension cells (Thermo Fisher Scientific) using the PEI precipitation method^80^. Recombinant human IgG1 antibodies were purified by affinity chromatography using Protein G Sepharose 4 Fast Flow (GE Healthcare), and dialyzed against PBS. Cv2F.4147, Cv2F.14107 and 76E1 variable domain genes were also cloned into human Igα1- and Fab-Igγ1-expressing vector^80^.Recombinant human IgA antibodies and Fab were purified by affinity peptide M-coupled agarose beads (Invivogen) and Ni Sepharose® Excel Resin (GE Healthcare), respectively.

#### ELISAs

ELISAs were performed as previously described^38,60^. Briefly, high-binding 96-well ELISA plates (Costar; Corning) were coated overnight with purified recombinant viral proteins (125 ng/well in PBS). After washings with 0.05% Tween 20-PBS (washing buffer), plates were blocked for 2 h with 2% BSA, 1 mM EDTA, 0.05% Tween 20-PBS (Blocking buffer), washed, and incubated with serially diluted human sera (1:100 and 3 consecutives 1:10 dilutions) or recombinant antibodies in PBS. For binding screening, anti-RBD antibodies were tested in triplicate at 1 µg/ml. Selected anti-RBD and anti-FP IgG antibodies were evaluated at 10 µg/ml followed by seven consecutives 1:4 and 1:10 dilutions in PBS, respectively. After washings, plates were revealed by incubation for 1 h with goat HRP-conjugated anti-human IgG (Jackson ImmunoReseach, 0.8 µg/ml final) and by adding 100 µl of HRP chromogenic substrate (ABTS solution, Euromedex) after washing steps. Optical densities were measured at 405nm (OD405nm), and background values given by incubation of PBS alone in coated wells were subtracted. Experiments were performed using HydroSpeed™ microplate washer and Sunrise™ microplate absorbance reader (Tecan Männedorf, Switzerland). For peptide-ELISA, anti-FP and control IgG antibodies were tested at 10 μg/ml and three consecutive 1:10 dilutions in 1% BSA, 0.1% Tween 20-PBS using the same procedure as previously described^38,60^. For the competition experiments, Wuhan RBD-coated plates were blocked, washed and then incubated for 2 h with biotinylated antibodies at a concentration of 25 ng/ml in the presence of IgG1 antibodies as potential competitors at 50 µg/ml, all performed in duplicate. After washings, plates were developed by adding HRP-conjugated streptavidin (1:2000 dilution, BD Biosciences) for 30 min and ABTS solution as described above. The percentage of competition was calculated as follows: 100 × [1- (OD^bio-IgG+comp^ / OD^bio-IgG^)].

#### Flow cytometry binding assay

Freestyle 293-F were transfected with pUNO1-Spike-dfur (Spike, SpikeV1 to V13 plasmids; Invivogen) and XBB.1.5, BA.2.86, EG.5.1 pHCMV expression vectors^78^ (1.2 μg plasmid DNA *per* 10^6^ cells) using the PEI-precipitation method. Forty-eight hours post-transfection, 0.3 × 10^6^ transfected and non-transfected control cells were incubated with IgG antibodies for 30 min at 4°C (1 μg/ml). For the assessment of anti-FP binding, transfected and non-transfected cells were incubated with anti-RBD and control antibodies (at 10 µg/ml) in presence the presence of Dy650 labeled-ACE2 or -FP antibodies - labeled using the Dylight 650 NHS ester kit (Thermo Scientific) - in 2 mM EDTA, 5% FBS-PBS for 2 h at room temperature. After washings, cells were incubated for 20 min at 4°C with AF647-conjugated goat anti-human IgG antibodies (1:1,000 dilution; Thermo Fisher Scientific) and LIVE/DEAD Fixable Viability dye Aqua (1:1,000 dilution; Thermo Fisher Scientific), washed and resuspended in PBS–paraformaldehyde 1% (Electron Microscopy Sciences). Data were acquired using a CytoFLEX flow cytometer (Beckman Coulter) and analyzed by FlowJo software (v10.8; FlowJo LLC). Fold increase in binding for anti-FP IgG antibodies was determined as follow: gMFI-APC^IgG + anti-RBD^ ^or ACE2^ / gMFI-APC^IgG alone^.

#### ACE2 binding inhibition assay

ELISA plates (Costar, Corning) were coated overnight with 250 ng/well of purified ACE2 ectodomain^38^, washed and blocked with ELISA blocking solution. For pre-Omicron variants, biotinylated tri-S (1 µg/ml in PBS) and RBD (2 µg/ml in PBS) proteins were incubated in the presence of antibodies at 2 µg/ml and 20 μg/ml, respectively. For Omicron variants, biotinylated RBD proteins (2 µg/ml in PBS) were incubated in the presence of antibodies at 200 μg/ml. After washing, plates were revealed by a 30 min-incubation with HRP-conjugated streptavidin (1:1000 dilution; BD Biosciences), followed by PBST washes and addition of ABTS solution. Percentage of inhibition (%) was calculated using the following formula: [1 - (OD^RBD+mAb^ / OD^RBD^)] x 100.

#### Viral-like particle neutralization assay

SARS-CoV-2 HiBiT-PsVLP Assay (Promega) was performed according to the manufacturer’s instructions. Briefly, SARS-CoV-2 HiBiT-PsVLPs mixed with antibodies (1 µg/ml for D614G and 10 µg/ml for BA.1 and BA.5) were incubated for 30 min at 37°C. Antibody-VLP complexes were then added to HEK293T LgBit cells in the presence of DrkBit Peptide followed by a 3 h incubation. Nano-Glo® Live Cell Reagent was added to the cells and after 15 min incubation, bioluminescence signals (RLU) were measured using a SAFAS Xenius plate reader (SAFAS Monaco). Percentage inhibition (%) was calculated as follows : 100 × [1 − (RLU^mAb^/ RLU^no mAb^].

#### SARS-CoV-2 S-Fuse neutralization assay

S-Fuse cells (1:1 mix of U2OS-ACE2-GFP1-10 and U2OS-ACE2-GFP11) were prepared and plated at a density of 2 x 104 per well in a μClear 96-well plate (Greiner Bio-One) as previously described^77^. SARS-CoV-2 virions (MOI 0.1) were incubated with antibodies at concentrations of 1 µg/ml for pre-Omicron variants and 10 µg/ml for Omicron variants in culture medium for 30 min at room temperature and then added to S-Fuse cells. For the most potent neutralizing antibodies, testing was performed at a starting concentration of 9 µg/ml, followed by 11 consecutive 1:4 serial dilutions. Cells were fixed 18 h later with 4% paraformaldehyde, washed, and stained with Hoechst dye (dilution 1:10,000; Invitrogen). Images were acquired with an Opera Phenix high-content confocal microscope (Perkin Elmer). The area of GFP-positive cells and the number of nuclei were quantified using with Harmony software 4.8 (Perkin Elmer). The neutralization % was calculated from the number of syncytia as follows: 100 × (1 - (value with IgG - value in “non-infected”) / (value in “no IgG” - value in “non-infected”)). IC_50_ values were calculated using Prism software (v.9.3.1, GraphPad Prism Inc.) by fitting replicate values using the four-parameters dose–response model (variable slope).

#### ADCP assay

PBMCs were isolated from healthy donors’ blood (Etablissement Français du Sang) using Ficoll Plaque Plus (GE Healthcare). Primary human monocytes were purified from PBMCs by MACS using Whole Blood CD14 MicroBeads (Miltenyi Biotech). Biotinylated Wuhan SARS-CoV-2 tri-S proteins were incubated with FluoSpheres™ NeutrAvidin™ beads (1 μm, yellow-green (505/515); Thermo Fisher Scientific; 20 μg of tri-S for 2 μl of beads), washed, and incubated for 30 min at room temperature. After PBS washes, tri-S-coupled beads 1:10000-diluted in DMEM were incubated for 1 h at 37°C with human IgG1 antibodies (at 5 μg/ml). The tri-S-bead-antibody mixtures were then incubated with 1×10^5^ human monocytes for 2 h at 37°C. 0.5% BSA, 2 mM EDTA, PBS solution, cells were fixed with 4% PFA–PBS and analyzed using a CytoFLEX flow cytometer (Beckman Coulter). ADCP experiments were analyzed using the FlowJo software (v10.8; FlowJo LLC). ADCP score corresponds to the percentage of FITC-positive cells.

#### ADCC assay

The ADCC activity of anti-RBD antibodies IgG antibodies was determined using the ADCC Reporter Bioassay (Promega) according to the manufacturer’s instructions. Briefly, 1×10^5^ Wuhan tri-S-transfected HEK293F cells were co-cultured with 6×10^4^ Jurkat-CD16-NFATrLuc cells in the presence or absence of anti-RBD or control mGO53 IgG antibody at 10 μg/ml. Luciferase was measured after 16 h of incubation using an GloMax Discoverer® plate reader (Promega). ADCC was measured as the fold induction of luciferase activity compared with the control antibody mGO53.

#### CD assay

The complement deposition (CD) of anti-RBD antibodies on Wuhan SARS-CoV-2 triS-expressing 293F cells was measured as previously described^84^. Briefly, 2×10^5^ triS-expressing 293F cells were cultivated in the presence of 50% normal or heat-inactivated human serum, and with or without IgG antibodies at 10 μg/ml. After 24 h, cells were washed with PBS and incubated for 30 min at 4°C with the live/dead fixable aqua dead cell marker (1:1,000 in PBS; Life Technologies) and PE-conjugated mouse anti-C3/C3b/iC3b IgG (2 µg/ml, 6C9; BD Biosciences) before fixation. Data were acquired using a CytoFLEX flow cytometer (Beckman Coulter) and analyzed using FlowJo software (v10.3; FlowJo LLC). CD scores were calculated by dividing the gMFI of live cells with mAb and normal serum by the gMFI of live cells without mAb.

### QUANTIFICATION AND STATISTICAL ANALYSIS

B-cell immunophenotyping and quantification of B-cell subset frequencies were performed using FlowJo software (v10.8; FlowJo LLC). Antibody classes were compared for functional data (neutralization, Fc-effector functions) using two-tailed Mann-Whitney test for total individual values and with Chi-square test with Yates’ correction for group frequencies. Statistical analyses were performed using GraphPad Prism software (v.8.2; GraphPad Prism Inc.). Heatmap representations were generated using GraphPad Prism or complex heatmap R package (v2.10.0; https://www.bioconductor.org/packages/release/bioc/html/ComplexHeatmap.html). PCA plots were generated using the factoextra package (v1.0.7; https://CRAN.Rproject.org/package=factoextra). The quantification of the synergistic effect and corresponding synergy map representations for antibody combinations was calculated using the SynergyFinder web application (v3.0) with the corrected Zero interaction potency (ZIP) model (https://synergyfinder.fimm.fi).

## French COVID Cohort Study Group

Marie Bartoli^1^, Alpha Diallo^1^, Soizic Le Mestre^1^, Christelle Paul^1^, Ventzislava Petrov-Sanchez^1^, Yazdan Yazdanpanah^1^, Cécile Ficko^2^, Catherine Chirouze^3^, Claire Andrejak^4^, Denis Malvy^5^, François Goehringer^6^, Patrick Rossignol^6^, Tristan Gigante^6^, Morgane Gilg^6^, Bénédicte Rossignol^6^, Manuel Etienne^7^, Marine Beluze^8^, Delphine Bachelet^9^, Krishna Bhavsar^9^, Lila Bouadma^9^, Minerva Cervantes-Gonzalez^9^, Anissa Chair^9^, Charlotte Charpentier^9^, Léo Chenard^9^, Camille Couffignal^9^, Marie-Pierre Debray^9^, Diane Descamps^9^, Xavier Duval^9^, Philippine Eloy^9^, Marina Esposito-Farese^9^, Aline-Marie Florence^9^, Jade Ghosn^9^, Isabelle Hoffmann^9^, Ouifiya Kafif^9^, Antoine Khalil^9^, Nadhem Lafhej^9^, Cédric Laouénan^9^, Samira Laribi^9^, Minh Le^9^, Quentin Le Hingrat^9^, Sophie Letrou^9^, France Mentré^9^, Gilles Peytavin^9^, Valentine Piquard^9^, Carine Roy^9^, Marion Schneider^9^, Richa Su^9^, Coralie Tardivon^9^, Jean-François Timsit^9^, Sarah Tubiana^9^, Benoît Visseaux^9^, Dominique Deplanque^10^, Jean-Sébastien Hulot^11^, Jean-Luc Diehl^11^, Olivier Picone^12^, François Angoulvant^13^, Amal Abrous^14^, Sandrine Couffin-Cadiergues^14^, Fernanda Dias Da Silva^14^, Hélène Esperou^14^, Ikram Houas^14^, Salma Jaafoura^14^, Aurélie Papadopoulos^14^, Alexandre Gaymard^15^, Bruno Lina^15^, Manuel Rosa-Calatrava^15^, Céline Dorival^16^, Jérémie Guedj^17^, Guillaume Lingas^17^, Nadège Neant^17^, Laurent Abel^18^, Victoria Manda^19^, Sylvie Behillil^20^, Vincent Enouf^20^, Yves Levy^21^ and Aurélie Wiedemann^21^

^1^Agence Nationale de Recherches sur le Sida et les Hépatites virales-Maladies Infectieuses Émergentes (ANRS-MIE), 75015 Paris, France

^2^Hôpital d’Instruction des Armées Bégin, Service des Maladies Infectieuses et Tropicales, 94160 Saint-Mandé, France

^3^Centre Hospitalier Régional Universitaire Jean Minjoz, 25056 Besançon, France

^4^Centre Hospitalier Universitaire, 80080 Amiens, France

^5^Centre Hospitalier Universitaire, 33000 Bordeaux, France ^6^Centre Hospitalier Régional Universitaire, 54000 Nancy, France ^7^Centre Hospitalier Universitaire, 76000 Rouen, France

^8^French Clinical Research Infrastructure Network (F-CRIN), 75013 Paris, France

^9^Hôpital Bichat, AP-HP, 75018 Paris, France

^10^Hopital Calmette, 59000 Lille, France

^11^Hôpital Européen Georges Pompidou, AP-HP, 75015 Paris, France

^12^Hôpital Louis Mourier, 92700 Colombes, France

^13^Hôpital Necker, AP-HP, 75015 Paris, France

^14^Institut National de la Santé et de la Recherche Médicale (INSERM), 75013 Paris, France

^15^INSERM UMR 1111, 69007 Lyon, France

^16^INSERM UMR 1136, 75646 Paris, France

^17^INSERM UMR 1137, 75018 Paris, France

^18^INSERM UMR 1163, 75015 Paris, France

^19^Hopital Lariboisière, AP-HP, 75010 Paris, France

^20^Institut Pasteur, Université Paris Cité, Molecular Genetics of RNA Viruses, F-75015 Paris, France

^21^Institut de Recherche Vaccinale (VRI), INSERM UMR 955, 94000 Créteil, France

## CORSER Study Group

Laurence Arowas^1^, Blanca Liliana Perlaza^1^, Louise Perrin de Facci^1^, Sophie Chaouche^1^, Linda Sangari^1^, Charlotte Renaudat^1^, Sandrine Fernandes Pellerin^2^, Cassandre van Platen^2^, Nathalie Jolly^2^, Lucie Kuhmel^3^, Valentine Garaud^3^, Hantaniaina Rafanoson^3^, Soazic Gardais^4^, Nathalie de Parseval^5^, Claire Dugast^5^, Caroline Jannet^5^, Sandrine Ropars^5^, Fanny Momboisse^5^, Isabelle Porteret^5^, Isabelle Cailleau^6^, Bruno Hoen^6^, Laura Tondeur^7^, Camille Besombes^7^, Arnaud Fontanet^7^

^1^ICAReB platform (Clinical Investigation & Access to Research Bioresources) of the Center for Translational Science, Institut Pasteur, F-75015 Paris, France

^2^Center for Translational Sciences, Institut Pasteur, F-75015 Paris, France

^3^Medical Center of the Institut Pasteur, Institut Pasteur, F-75015 Paris, France

^4^G5 Infectious Disease Epidemiology, Institut Pasteur, F-75015 Paris, France

^5^Institut Pasteur, F-75015 Paris, France

^6^Direction de la recherche médicale, Institut Pasteur, F-75015 Paris, France

^7^Emerging Diseases Epidemiology Unit, Institut Pasteur, F-75015 Paris, France

